# “Optimality of motor and clutch mechanical properties in a generalized model for cell traction forces”

**DOI:** 10.1101/2023.02.20.529282

**Authors:** Roberto Alonso-Matilla, Paolo P. Provenzano, David J. Odde

## Abstract

The mechanical properties of the extracellular environment influence cell behavior, such as cell morphogenesis, cell migration and cell fate. During the cell migration cycle, cells extend plasma membrane protrusions via actin polymerization, actin filaments move retrogradely via myosin motors and actin polymerization, and the formation of adhesion complexes commonly called clutches mediate the production of traction forces on the extracellular microenvironment allowing the cell to move forward. In this study, we revisited the motor-clutch model for cell migration and derived its mean-field counterpart to study force production of cellular protrusions on compliant substrates. We derived an analytical expression for the optimum substrate stiffness that maximizes force transmission, without unknown adjustable parameters that is valid for all motor-clutch ratio regimes; and determined the required conditions for the protrusion to operate in the clutch-dominated “stalled” regime, the motor-clutch “balanced” regime and motor-dominated “free flowing” regime. We explored these regimes theoretically and obtained an analytical expression for the traction force produced in the limits of low and high myosin activity. We discovered the existence of an optimum clutch stiffness for maximal cellular traction force and identified a biphasic dependence of traction force produced by protrusions on rigid substrates on motor activity and unloaded myosin motor velocity, results that could be tested via novel molecular tension sensor experiments. We additionally showed that clutch reinforcement shifts the optimum substrate and clutch stiffnesses to larger values whereas the optimum substrate stiffness is found to be insensitive to changes in clutch catch bond properties. Overall, our work reveals that altering cell adhesion levels, actomyosin activity and rigidity of clutches and extracellular matrix can cause nontrivial traction force changes and modulated cell migration capabilities that provide insights to effectively design novel cell and stromal-based therapies to treat development and human diseases, such as cancer invasion and metastasis.

**STATEMENT OF SIGNIFICANCE:** Adherent cells produce mechanical forces on their environment that critically regulate cell adhesion, signaling and function, essential during developmental events and human disease. Despite recent progress, cell-generated force measurements across a wide range of extracellular stiffnesses and cell states have faced numerous technical challenges. Our mean-field study provides a new generalized analysis of regulation of force transmission by modulation of cellular components and extracellular rigidity. We identify the existence of an intermediate stiffness of cell adhesion complexes for maximum force transmission and find that effective force transmission on rigid environments reduces to a competition between cell adhesion reinforcement and load-dependent dissociation kinetics. The developed model provides important insights to aid design of novel therapeutic strategies for cancers and other diseases.

## INTRODUCTION

Cell migration plays a pivotal role in many biological processes, such as embryonic morphogenesis, wound healing, and cancer progression. A grand challenge in cell biology is to understand how cells within an organism migrate through different environments. During migration, cells undergo a complex series of events that occur in a highly dynamic and periodic fashion (1,2). According to the motor-clutch hypothesis (3), cells start off the migration cycle by the extension of cell membrane protrusions driven by actin polymerization, followed by the formation of complex adhesion structures and generation of traction forces (4). Adhesion complexes function as molecular clutches by mechanically coupling the actin cytoskeleton to the extracellular substrate via membrane-bound receptors. Clutches cooperatively resist the forces arising from the rearward flow of actin filaments and shunt it to the substrate, allowing the cell to move forward (5–7). The flow of actin away from the leading edge, known as retrograde flow, emerges as actin filaments are subject to rearward forces produced by the leading-edge membrane due to actin polymerization (8) as well as by myosin motors as they bind and pull filaments away from the leading edge (9). Actin polymerization, adhesion formation and myosin forces are therefore mechanically coordinated allowing cells to migrate and explore their extracellular environment.

How cells sense and response to the mechanical features of their environment has been the focus of many studies (10–14). According to the motor-clutch model, coupling between the extracellular substrate and the actin cytoskeleton by molecular clutches permits the transmission of actomyosin-generated forces to the substrate, allowing cells to feel the mechanical properties of the environment (15). The motor clutch model exhibits rigidity sensing, that emerges from load-dependent clutch-bond dissociation rates and load-dependent myosin motor force generation (2,10,16,17), and captures the reported biphasic dependence of cell migration on substrate adhesion strength, first postulated by DiMilla et al. (18) and later confirmed experimentally (19,20). Cell motility (12,20–26) and traction forces (12,13) have been reported to exhibit a biphasic dependence on substrate adhesivity/stiffness. Also, some cells produce higher traction forces on stiffer substrates (27–30), whereas other cells produce higher traction forces on softer substrates (2). Together, these studies suggest the existence of an optimal substrate stiffness for maximal traction force.

Previous mean-field model studies addressed the production of traction forces on infinitely rigid substrates (31–33), and in compliant linearly elastic (16) and viscoelastic (34) substrates. In this study, we introduce a mean-field representation of the motor-clutch model (2) with the aim of gaining a better theoretical understanding of traction force production of individual cellular protrusions on elastic compliant substrates. Particularly, we apply scaling analysis to our model with the purpose of deriving an analytical expression for the optimum substrate stiffness, i.e. the substrate stiffness that maximizes traction forces. An expression for the optimum substrate stiffness was previously derived in (16), where the probability density function of bound clutch forces was assumed to obey a gamma distribution. The final expression for the optimum stiffness was only valid for protrusions with balanced number of motors and clutches and was left as a function of an unknown parameter ε, the fraction of the theoretical maximum load that a protrusion can generate. In this work, we relax the assumption made in (16) and derive a more general expression for the optimum substrate stiffness without an unknown adjustable parameter and it is applicable to all motor-clutch ratio regimes. We find that the optimum stiffness is not sensitive to clutch stiffness and corresponds to the situation where the characteristic time for available clutches to mechanically link actin filaments to the extracellular substrate equals the characteristic time for clutches to rupture due to force. We find that motor-dominated protrusions are characterized by a myosin-independent optimum stiffness, clutch-dominated protrusions are characterized by stall conditions and stiffness-independent traction forces (10,16), and balanced clutch-motor protrusions are characterized by an optimum stiffness sensitive to motor activity. We explore the three different cellular types in detail, derive analytical expressions that indicate the required conditions for the cell to operate at each cellular regime, and obtain analytical expressions for the traction force generated by cellular protrusions on non-compliant substrates, in the limits of low and high myosin activity. Our mean-field model results allow us to recognize a strong dependence of force transmission on clutch stiffness, particularly for protrusions on low compliant substrates, and to identify the existence of an optimum clutch stiffness for maximal cellular traction force, an optimum that is nearly insensitive to substrate stiffness and that strongly depends on clutch kinetics. In addition, we capture a biphasic dependence of force transmission on motor activity on rigid substrates and identify an intermediate myosin load-free velocity parameter for maximum traction force. Finally, we explore the dependence of traction force generation on clutch reinforcement and catch bond dynamics. Whereas clutch reinforcement can significantly shift the optimum substrate and clutch stiffnesses to larger values, increasing substantially the traction force produced by protrusions that display load-and-fail dynamics, catch bond properties can modulate force transmission but do not affect the optimum substrate stiffness for maximum force. We found that force transmission on rigid substrates can be rescued by clutch reinforcement provided that it is sufficiently strong to prevent frictional slippage. We numerically solve the mean-field model equations and find perfect agreement between results from the mean-field model and the stochastic model (2). Our mean-field approach additionally allowed us to carry out an in-depth exploration of the distribution of clutch force states during the loading cycle.

## MODEL DESCRIPTION

In this section, we introduce a mean-field representation of the motor-clutch model to study the dynamics of force transmission at individual cellular protrusions. According to the motor-clutch hypothesis, myosin motors bind and pull actin filaments retrogradely away from the leading edge producing retrograde forces. Forces on actin filaments are transmitted to the extracellular matrix through protein complexes, commonly called clutches, that mediate the production of traction forces on the surrounding compliant substrate allowing the cell body to propel forward. Molecular clutches bind — that is, couple actin filaments to the compliant substrate, with vanishing extension at a force-independent rate k_on_. Clutches form slip bonds and unbind with an effective dissociation rate that increases exponentially with force according to Bell’s law 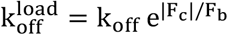 (35), where k_off_ is the unloaded clutch dissociation rate, F_b_ is the characteristic bond rupture force, and F_c_ is the force on the clutch. The simplest motor-clutch description is to consider molecular clutches and surrounding substrate as Hookean elastic materials. Therefore, the clutch force is given by Hooke’s law F_c_ = κ_c_x_c_, where κ_c_ is the effective clutch stiffness and x_c_ is the clutch extension, measured with respect to its resting value. To study clutch elongation dynamics, we introduce a probability density function P_b_(x_c_, t) of finding a bound clutch with elongation x_c_ at time t. Conservation of P_b_ implies (see Supplemental Information):

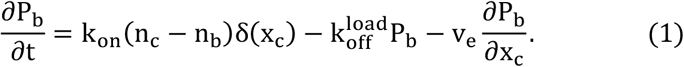

The first term on the RHS of Eq. (1) corresponds to the rate of change of the probability density due to clutch binding kinetics, where δ(x_c_) is the Dirac delta function which enforces clutches to engage in an unloaded configuration. The number of bound clutches that mechanically connect the cytoskeletal filaments with the substrate is denoted by n_b_(t), and n_c_ is the total number of available clutches in the cellular protrusion. The second term on the RHS of Eq. (1) is the rate of change of the probability density due to clutch dissociation kinetics, and the last term of Eq. (1) accounts for the rate of change of the probability density due to force-mediated clutch extension, where the clutch elongation rate is v_e_ = v_act_ – dx_s_⁄dt, where v_act_ is the actin filament velocity and dx_s_/dt is the substrate deformation rate. We follow therefore the standard motor-clutch view and assume that all bound clutches equally deform at any time, where the actin binding domains of clutches move at the actin retrograde flow velocity v_act_ and their substrate binding domains move rigidly with the substrate. Myosin motors obey a linear force-velocity relation; thus, the total force exerted by myosin motors on the actin filament bundle is F_act_ = n_m_F_m_(1 – v_act_⁄v_u_), where n_m_ is the total number of myosin motors in the protrusion, F_m_ is the stall force exerted by one myosin motor and v_u_ is the load-free velocity of myosin motors. For convenience, we define the first two moments of the probability density over all possible clutch extensions: 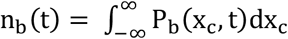 and 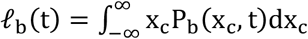, where n_b_ is the above-mentioned number of bound clutches, and 𝓁_b_ is the sum of clutch extensions. We have defined P_b_(x_c_, t) on an infinite domain of clutch extensions to guarantee numerical stability. A negative clutch extension means that the substrate binding domain of the clutch falls behind of its actin binding domain, a phenomenon that we rarely expect to occur in physiological conditions. Governing equations for the first two moments can be obtained from simple quadrature of the conservation equation for the probability density function, yielding 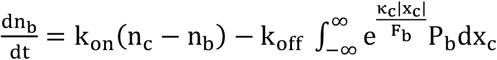, and 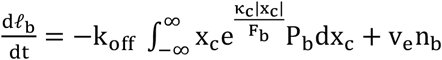. A force balance on the substrate allows us to calculate the traction force exerted on the substrate T as a function of the first-order moment of the probability density: T = κ_s_x_s_ = κ_c_𝓁_b_, where κ_s_ is the substrate stiffness constant and x_s_ is the substrate deformation. We take the time-derivative of the traction force and use Eqs. (2,3) to obtain an expression for the clutch elongation rate

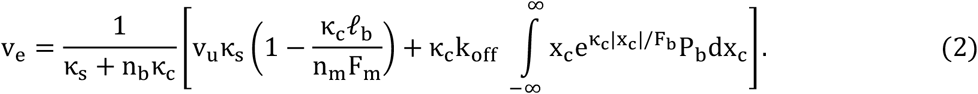

For convenience we use dimensionless equations where we scale variables using 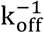 as time scale, n_m_F_m_/n_c_κ_c_ as length scale, n_m_F_m_k_off_/n_c_κ_c_ as velocity scale, and n_c_F_b_ as force scale: 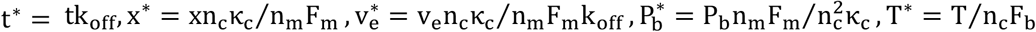, where we have additionally normalized the probability density function by the total number of clutches n_c_. The fraction of bound clutches is denoted as 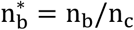. Henceforth, any variable accompanied by an asterisk is dimensionless. Nondimensionalization of the governing equations yields four dimensionless groups: F = n_m_F_m_⁄n_c_F_b_, τ = k_on_⁄k_off_, ω = v_u_κ_c_⁄F_b_k_off_, K = κ_s_⁄n_c_κ_c_. The myosin activity parameter F is the ratio of the total myosin stall force n_m_F_m_ over the characteristic maximum clutch elastic force n_c_F_b_. The clutch kinetic parameter τ is the ratio of the clutch association constant k_on_ over the clutch unloaded dissociation constant k_off_. The parameter ω represents the ratio of the characteristic clutch unloaded dissociation time over the characteristic clutch loaded dissociation time. The parameter ω can also be described as the dimensionless myosin load-free velocity, or as the ratio of the characteristic elongation that a single clutch undergoes in the absence of cooperative clutch effects over the characteristic clutch rupture length. Notice that only large ω values are physiologically relevant (ω ≫ 1). Finally, the parameter K is the ratio of the substrate stiffness κ_s_ over the maximum effective clutch stiffness n_c_κ_c_. With these scalings, the conservation equation (1) and dimensionless clutch strain rate 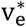 read

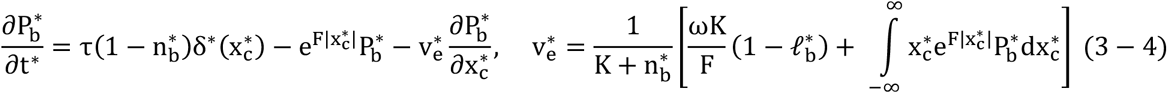

where the first moment of the probability density over clutch extensions takes the form 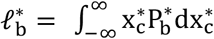. Notice that the traction force exerted on the compliant substrate is 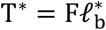. Throughout the manuscript, we will use an overline symbol acting on a variable to denote the long time-averaged value of the variable.

The developed mean-field theory is an extension of the model by Bangasser & Odde (16), where a probability that each existing clutch in the protrusion is mechanically linking the actin cytoskeleton and the extracellular substrate was introduced. In this work, we introduce a multibond probability density of finding any clutch with a given extension at any instantaneous time and derive its conservation equation (see Supplemental Information). In particular, our approach relaxes the previous assumption that motors and clutches are balanced, without requiring an unknown parameter to define the optimum, and analytically investigates previously uncharacterized optimality of clutch stiffness, catch bond behavior, and bond reinforcement behavior, the latter two of which are common features of motor-clutch systems experimentally. Thus, the present treatment allowed us to make a more complete and generalized analysis of cellular force transmission, and further explored the entire motor-clutch model parameter space.

## RESULTS

### Production of traction forces on individual cellular protrusions exhibit three different regimes: a motor-dominated regime, a motor-clutch balanced regime and a clutch-dominated stalled regime

Prior motor-clutch studies (2,10,16) identified the existence of three different traction force production regimes: a stalled regime characterized by clutch-dominated protrusions, a balanced regime characterized by a motor-clutch balanced protrusion and a free-flowing regime characterized by motor-dominated protrusions. Our mean-field model helped us identify and derive analytical expressions for the three key timescales that control traction force production in the motor-clutch model: the characteristic time for available clutches to bind (i.e. mechanically couple actin filaments to the extracellular substrate) t_bind_, the characteristic time for clutches to rupture due to force t_rupt_, and the characteristic time for the substrate to reach its maximal deformation 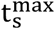. These three timescales depend on the different parameters of the model, dictate the cellular regime at which the protrusion operates, and are essential to understand traction force production and optimality conditions. We numerically solve the governing equations of the mean-field model (Eqs. (3) and (4)) using a finite-difference algorithm (see Supplemental Information) and find a very good agreement with the stochastic motor-clutch model version (2). We explore the three different cellular regimes in detail below.

#### Clutch-dominated stalled regime

A distinct feature of the clutch-dominated regime, previously identified as the stall regime (10,16), is that the protrusion operates at its maximum efficiency by producing the maximum possible traction force, i.e. the total myosin stall force 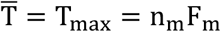 (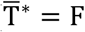 in dimensionless form), throughout a wide range of substrate stiffnesses; hence no single optimum substrate stiffness for maximum traction force exists. The clutch-dominated regime is therefore characterized by stiffness-independent traction forces as well as stiffness independent binding/unbinding kinetics, as shown in Fig. (1) by the mean-field numerical solutions and stochastic motor-clutch model solutions of the time-averaged traction force (Fig. 1A) and time-averaged fraction of bound clutches (Fig. 1B, inset). The mean-field model captures the time-averaged traction force dynamics. In an unloaded protrusion, actin filaments flow rearwards at the myosin load-free velocity. The first clutches that connect the fast-moving actin filaments with the deformable substrate undergo large extensions, since there is not enough elastic resistance against myosin pulling forces. This is shown in Fig. (1B) by the fat tail of the probability density function at short times. As time goes by, more clutches mechanically couple the actin cytoskeleton with the compliant substrate, as shown in Fig. (1B, inset) and manifested by the rise of the area under the probability density curve in Fig. (1B), and elastic energy builds up in both substrate and bound clutches, that slows down actin retrograde flows. Low actin filament velocities lead clutches to dissociate stochastically before they can reach very large elongations, as demonstrated in Fig. (1B) by a reduction in the skewness of the probability distribution function at longer times. During loading in strong-clutch protrusions, clutch binding dominates clutch unbinding, and no clutch failure avalanche occurs at any instant in time, as there are always enough bound clutches resisting myosin pulling forces. The absence of load-and-fail dynamics is confirmed in Figs. (1A), and (1B); and occurs for cellular protrusions that strain the substrate to its maximum deformation before clutch bonds break due to force 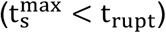— that is, F-actin retrograde flows vanish and the protrusion stalls. This inequality implies (see derivation below) that protrusions with a motor activity parameter lower than a threshold 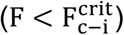 lie on the clutch-dominated regime and produce maximum available traction forces. Numerical values for 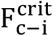 are displayed in Fig. (1C) as a function of τ and ω for both soft and rigid substrates, and an analytical expression for the critical activity parameter 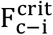 has been estimated for soft substrates (see derivation below). The critical parameter 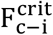 monotonically increases as τ increases, as higher myosin forces are required for clutch failure avalanches to occur in high-clutch-binding-rate protrusions. The dependence of 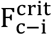 on the dimensionless load-free velocity ω is less straightforward. Stronger retrograde flows exist at the beginning of loading for larger values of ω. On very soft substrates, loading is so slow that the initial strong retrograde flows mainly deform the substrate, barely affecting clutch dynamics at long times. Accordingly, 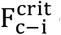 on soft substrates is nearly independent of ω, as shown in Fig. (1C). On rigid substrates, on the other hand, clutches build up elastic energy very fast; thus, higher values of ω enhance clutch deformations at the onset of the loading cycle favoring load-and-fail dynamics over stall conditions. As a result, lower myosin forces are required for load-and-fail dynamics to occur on high-ω protrusions— that is, 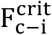 on soft substrates is nearly independentincreasing ω on rigid substrates, as shown in Fig. (1C).

**FIGURE 1.**
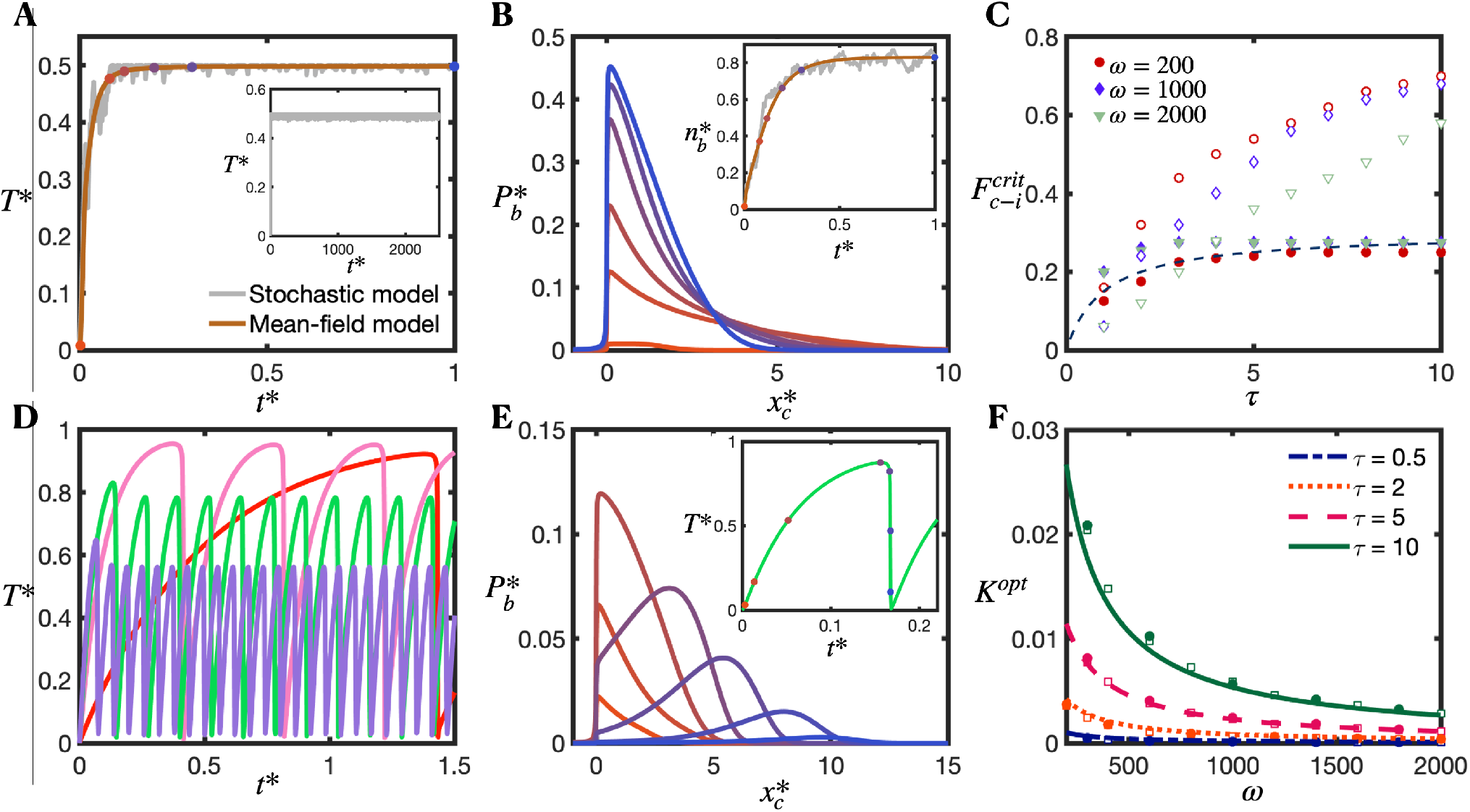
Clutch-dominated protrusions produce maximum possible traction forces and do not exhibit clutch failure avalanches whereas cellular protrusions that operate in the intermediate regime features load-and-fail dynamics and produce maximum traction forces at an intermediate substrate stiffness. Temporal evolution of the dimensionless traction force (A), probability density function (B) and fraction of bound clutches (B, inset) in a clutch-dominated protrusion. The dimensionless probability density function is obtained by the mean-field model and sampled at the time-points marked by symbols in (A, B inset). Color schemes of symbols in (A, B inset) and lines in (B) correspond to the same time-points. In the clutch-dominated regime, long-time traction force and fraction of bound clutches approach the total myosin stall force T = n_/_F_/_ and unloaded equilibrium fraction of bound clutches n_b_ = n_c_k_on_/(k_on_ + k_off_), respectively. Brown: mean-field model solution, grey: a single-trajectory of the stochastic model. A, inset: long-time stochastic single-trajectory. Parameter values: F = 0.5, τ = 10, ω = 500, K = 1. (C) Critical activity parameter 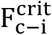 that sets the border between the clutch-dominated regime and the intermediate regime, as a function of the clutch kinetic parameter τ, for three values of the dimensionless myosin load-free velocity ω. Protrusions with 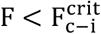 belong to the clutch-dominated regime and produce maximum available traction forces. Solid and open symbols correspond to the critical activity parameter in soft and rigid substrates, respectively. The dashed line is our analytical solution for soft substrates obtained via scaling analysis (Eq. 5). (D) Time-evolution of dimensionless traction force for four different substrate stiffnesses: K = 0.001 (red), K = 0.005 (pink), K = 0.008 (green) and K = 0.01 (purple). Parameter values: F = 1, τ = 10, ω = 2000. (E) Temporal evolution of the dimensionless probability density function 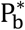 and dimensionless traction force (inset). Symbols and color scheme in inset correspond to the time points where 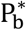 is sampled. Parameter values: F = 1, τ = 10, ω = 2000, K = 0.005. (F) Dimensionless optimum substrate stiffness 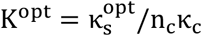 as a function of the dimensionless myosin load-free velocity ω, for four different values of the clutch kinetic parameter τ. The dimensionless clutch binding time scales as 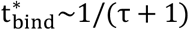, whereas the dimensionless clutch rupture time scales as 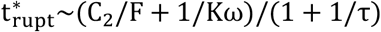. Solid lines correspond to our derived analytical solution (Eq. 6), open symbols correspond to the numerical solution of the mean-field model (Eqs. (3,4)), and closed symbols correspond to the numerical solution of the stochastic model. Parameter value: F = 2.

#### Motor-clutch balanced regime

As its name suggests, the intermediate regime is a transitional regime between the free-flowing motor-dominated regime and the clutch-dominated stalled regime, where motors and clutches balance each other. The intermediate regime is characterized by load-and-fail dynamics and by the existence of a motor-sensitive optimum substrate stiffness for maximum traction force, as depicted in Fig. (S1) for F∼1. Cycles of loading and failure are illustrated in Figs. (1D) and (S2) for different substrate stiffnesses. The mean frequency between two consecutive clutch cascading failure events decreases with substrate compliance, as shown in Fig. (1D), and in agreement with Chan & Odde (2). At the beginning of loading, the elastic energy built up in the system is undertaken by the substrate/clutches on soft/stiff environments, respectively. Therefore, clutch lifetimes depend inversely on substrate rigidity, and higher frequency of clutch failures will be observed on more rigid environments, as shown in Fig. (S1C). In the beginning of the loading cycle, the number of bound clutches is independent of substrate stiffness, being mainly governed by binding kinetics, as Fig. (S2B) shows. During this early phase, protrusions produce larger traction forces on more rigid substrates (Fig. 1D)). Later in the loading cycle, some clutches cannot sustain load any longer and eventually their clutch bonds break. The net clutch unbinding rate becomes greater than the clutch binding rate and the number of bound clutches begins to drop (Fig. (S2B)), leading to an increase in clutch elongation rate (Fig. (S2C)) and a reduction in substrate deformation rate (Fig. (1D)). As clutches gradually dissociate, the total load is redistributed among the remaining bound clutches, with an increase of load per clutch (Fig. (1E)). Eventually, the load cannot be sustained by the remaining clutches, and an instability occurs that eventuates in the rupture of all clutch bonds, a sudden fall in traction force (2), and the beginning of a new loading cycle. Notice that the duration of the clutch unbinding phase (dn_b_/dt < 0) is much longer than that of the clutch binding phase (dn_b_/dt > 0), especially on very soft substrates.

The boundary between the clutch-dominated stalled regime and the balanced regime is set by the condition 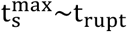 — that is, the time required to reach stall conditions matches the time needed for clutch bonds to break due to force. We perform some scaling analysis (see Supplemental Information) and find that the protrusion operates in the clutch-dominated regime when

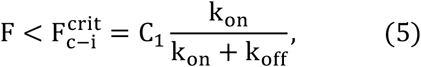

where C_1_ = 0.4 is a constant that we estimate by fitting Eq. (5) with our numerical results. The protrusion belongs to the clutch-dominated regime and produces the maximum possible traction force when the ratio between the total myosin stall force n_m_F_m_ and the equilibrium clutch elastic force n_c_F_b_k_on_/(k_on_ + k_off_) is lower than a constant of order 1. Figure (1C) shows a very good agreement between our theoretical formula and numerical results.

#### Motor-dominated regime

This regime is characterized by a myosin-independent optimum stiffness. In this regime, we find that the cell can operate under motor-dominated conditions and still sense the rigidity of the environment (see Fig. S1A, F = 10 and F = 20). In strong-motor protrusions, any changes in motor activity will not appreciably affect the dynamics of the protrusion (i.e. traction forces, clutch binding/unbinding kinetics, etc.). This is shown in Fig. S1A, where curves for F = 10, F = 20 and F = 100 (not shown for clarity) lie on top of each other and reach a maximum at an intermediate substrate stiffness. Before any clutch links the actin cytoskeleton with the substrate, actin filaments flow rearwards at the myosin load-free velocity. Once an individual clutch binds, it provides elastic resistance. In the high-myosin limit, both clutch elongation and substrate elongation rates are myosin-independent, as demonstrated in our simple derivation (see Supplemental Information), which indicates that motor-dominated protrusion dynamics are myosin insensitive. In the next section, we show that protrusions belong to the strong-motor regime when 0.05 k_on_n_c_F_b_/(k_off_ n_m_F_m_) ≪ 1. Motor-dominated protrusions also undergo load-and-fail dynamics. Higher myosin forces strengthen retrograde flows causing a reduction in the lifetime of clutch bonds, and provoking more frequent clutch rupture avalanches, as observed in Fig. (S1C), and in agreement with previous studies (2). The previously identified free-flowing cellular state (10,16) is a particular case of the motor-dominated regime for motor-dominated protrusions on sufficiently rigid substrates. Sufficiently rigid substrates build up traction forces at a fast rate, clutches reach a high tensional state very rapidly and a state of frictional slippage takes place where clutches continually dissociate by force before additional clutches can associate and share the load with the already bound clutches, limiting traction force production.

### Optimum substrate stiffness for maximum traction force is reached when the clutch binding time equals the clutch rupture time

In this section we apply scaling analysis to the mean-field model with the purpose of deriving an analytical expression for the optimum substrate stiffness, i.e. the substrate stiffness that maximizes traction forces. When clutch bonds in the protrusion break before stall conditions are reached, i.e. 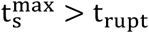, the protrusion undergoes periods of clutch loading and unloading (load-and-fail behavior). If the substrate is too rigid, the characteristic time for clutches to bind t_bind_ is much larger than the characteristic time for clutches to rupture due to load t_rupt_ (t_bind_ ≫ t_rupt_), and clutch bonds break before they have enough time to form large stable adhesions. An early clutch failure cascade thus results in traction forces that are far below their optimum values. If the substrate is too soft, t_bind_ ≪ t_rupt_, most of available clutches mechanically link the actin cytoskeleton with the substrate long before the clutch bonds break by force. Yielding substrates undergo high strain rates that lower clutch deformation rates, and the protrusion spends most of the clutch loading cycle in a state of low traction force production and high retrograde flow, leading to force transmission far below its optimum value. Therefore, there must exist an intermediate substrate stiffness that maximizes mean traction force production (10,16). Previously, we found that optimum conditions for maximum traction force are reached when the time needed for available clutches to bind equals the cycle time (10,16). Here, we identify the cycle time as the characteristic time for clutches to rupture due to load, thus we hypothesize that optimum conditions are reached when t_bind_∼t_rupt_. We apply scaling analysis to estimate t_bind_ and t_rupt_ (see Supplemental Information); we get:

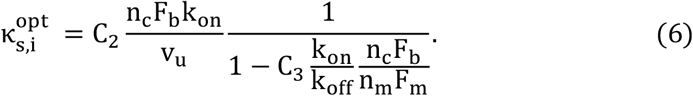

Our scaling analysis allows us to obtain the optimum substrate stiffness up to two unknown constants C_2_ and C_3_. We estimate the constants by fitting Eq. (6) to our numerical results. We get C_2_ = 0.4, C_3_ = 0.05. Figures (1F) and (S3) show that our theoretical expression is in very good agreement with our numerical results for all the motor-clutch dimensionless parameters. The optimum substrate stiffness in a protrusion that belongs to the intermediate regime, 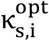, is myosin sensitive and independent of the effective clutch stiffness κ_c_ as previously reported in other studies (10,16). This apparent counterintuitive result can be explained by realizing that both characteristic clutch rupture length 𝓁_rupt_ and the characteristic clutch extension rate scale inversely to clutch stiffness. This implies that clutch rupture time is independent of clutch stiffness. Since neither the clutch binding rate nor the clutch rupture time depend on clutch stiffness, the substrate stiffness that maximizes traction forces does not depend on clutch stiffness, as Eq. (6) shows. We expect this result to hold for all physiologically relevant parameter values. In the non-physical situation where clutches dissociate stochastically instead of by force (small ω), the characteristic clutch rupture length would not be F_b_⁄κ_c_ anymore, and optimum substrate stiffness would *a priori* depend on clutch stiffness. We have assumed that clutch dissociation rates increase exponentially by force according to Bell’s law. Protrusions with clutches that have a different lifetime-extension dependence will, in principle, operate at optimum substrate stiffnesses that depend on clutch stiffness. The optimum stiffness in motor-dominated protrusions reduces to 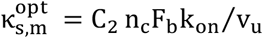, where we have taken the limit C_3_k_on_n_c_F_b_/k_off_n_m_F_m_ → 0. We also find very good agreement between our theoretical prediction, the numerical solution of the mean-field model and the solution of the dimensionless version of the stochastic motor-clutch model (2) for motor-dominated protrusions, as shown in Fig. (S3, right). According to our derived expression, the optimum substrate stiffness is proportional to the characteristic maximum clutch elastic force n_c_F_b_ and inversely proportional to the distance that unloaded actin filaments translocate within the characteristic clutch binding time v_u_/k_on_. Our theoretical solution also suggests that in the high-motor regime, the optimum substrate stiffness is myosin-insensitive.

### Traction force production is maximal at intermediate clutch stiffnesses

In the previous sections we showed that the optimum substrate stiffness 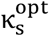 is insensitive to changes in clutch stiffness, κ_c_. Force transmission, however, could *a priori* depend on clutch stiffness when the substrate stiffness differs from the optimum. We now proceed to explore the dependence of force transmission on clutch stiffness. To investigate this effect, we notice that among the four dimensionless parameters introduced, only two depend on κ_c_: the parameter ω that is proportional to κ_c_, and the dimensionless substrate stiffness K that depends inversely on κ_c_. Accordingly, we introduce a new variable, the clutch stiffness parameter β_c_, such that *ω* = *ω*′β_c_ and *K* = *K*′/β_c_, where *ω*′ and *K*′ are, respectively, the values of the parameter *ω* and the dimensionless substrate stiffness *K* when β_c_ is equal to 1. Consequently, by definition, increasing the value of the clutch stiffness parameter β_c_ is equivalent to proportionally increasing the effective clutch stiffness κ_c_. We investigated the dependence of the time-averaged force transmission on clutch stiffness by numerically solving our mean field model equations. Our numerical results identify the existence of an optimum clutch stiffness for maximum traction forces (Figs. 2A and 2B). Near-perfect agreement is found between the mean field model solutions and the stochastic solutions. In the stiff-clutch limit, clutches provide enhanced elastic resistance to deformations, decreasing their strain rates. Their characteristic clutch rupture length, however, is inversely proportional to their stiffness (assuming that the clutch rupture force is constant). In the stiff-clutch limit, clutch lifetimes are low, especially on non-compliant substrates, and this is reflected in a lower mean number of bound clutches, as shown in Fig. (S4). Even though the traction force buildup is extremely fast at the beginning of loading, the strong dependence of clutch lifetimes on their stiffness becomes the dominant effect and the resulting traction forces transmitted to the substrate are also low. In the soft-clutch limit, clutch extension rates are higher, but clutches disengage before high traction forces are produced, since the soft-clutch limit is associated with a slow build-up of traction forces, as shown in Fig. (2C). Consequently, there exists an optimum clutch stiffness that maximizes force transmission. We proceed to quantify whether the clutch stiffness optimum could be relevant in physiological conditions. Under the assumption that v_u_ = 120nm ∙ s^−1^, F_b_ = 2pN, k_off_ = 1s^−1^ and τ = 10 (k_on_ = 10s^−1^), the mean-field model predicts that the optimum clutch stiffness is 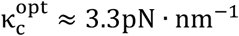, a clutch stiffness that is within the range of physiological clutch stiffness values. Interestingly the sensitivity of force transmission on clutch stiffness is significantly higher for protrusions on rigid substrates than for protrusions on soft substrates, as shown in Figs. (2A) and (2B). This can be explained by noticing that, on stiff substrates, the load-and-fail dynamics are largely governed by the mechanical properties of clutches. Consequently, the sensitivity of traction forces to changes in clutch stiffness is expected to be higher on this type of substrates. The optimum clutch stiffness increases for higher values of the clutch kinetic parameter τ, as shown by the shift in the optimum towards the right in Figs. (2A) and (2B) and decreases with increasing values of the myosin activity parameter *F*, as shown in Fig. (2D). Faster clutch recruitment or lower myosin activity strengthens clutches over motors, and stiffer clutches will accelerate the build-up of traction forces, increasing the overall force transmission. This is consistent with Eq. (13), where we show in the next section that the production of traction forces on rigid substrates in the limit of low myosin activity increases as the clutch stiffness increases. Direct comparison of Figs. (2A) and (2B) indicates that the optimum clutch stiffness is barely sensitive to substrate compliance.

**FIGURE 2.**
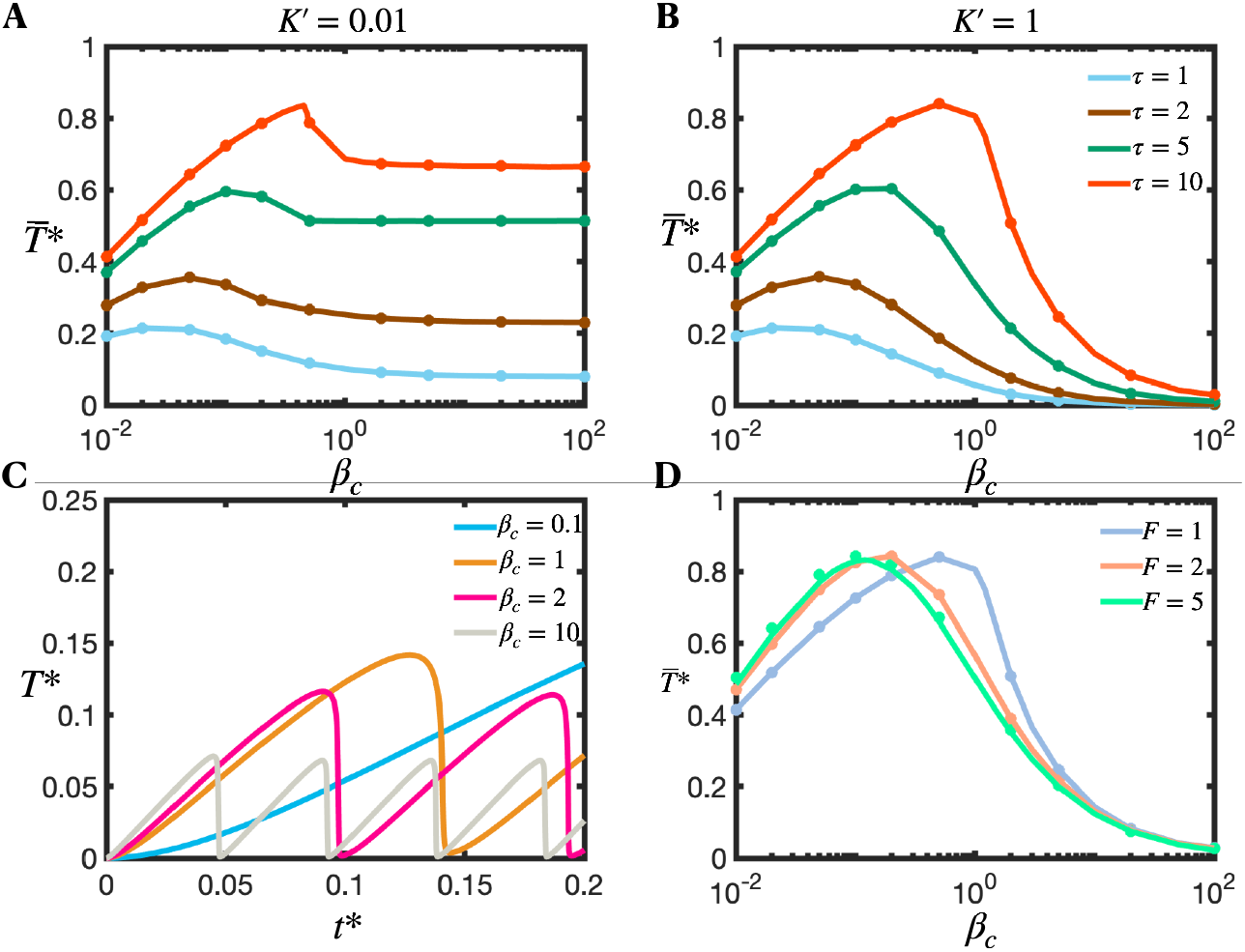
Traction force transmission is maximum at intermediate values of clutch stiffness— (A,B) Dimensionless time-averaged traction force 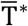 as a function of the clutch stiffness parameter β_c_ for different values of the clutch kinetic parameter τ and for two values of the substrate stiffness parameter *K*′ (*K* = *K*′/β_c_): (A) *K*′ = 0.01 and (B) *K*′ = 1. Cellular traction forces on rigid substrates are very sensitive to clutch stiffness. The optimum clutch stiffness (optimum β_c_) increases with the value of the clutch kinetic parameter. Parameter values: *F* = 1, *ω*′ = 200 (*ω* = *ω*′β_c_). Notice that κ_c_ ∝ β_c_. (C) Time-evolution of traction force for four different values of the clutch stiffness parameter. Parameter values: *F* = 1, *ω*^M^ = 200, *K*^M^ = 0.01, *τ* = 1. (D) Dimensionless time-averaged traction force as a function of the effective clutch stiffness parameter for three different values of the myosin activity parameter F. Parameter values: *ω*′ = 200, *K*′ = 1, τ = 10. Solid lines correspond to the numerical solution of the mean-field model (Eqs. (3) and (4)), and symbols correspond to the numerical solution of the stochastic motor-clutch model.

### Traction forces produced by protrusions on rigid substrates reach a maximum at intermediate myosin levels and unloaded myosin velocities

In the previous sections, we used two approaches to study the dynamics of cell protrusions: a stochastic Langevin-type approach (2) and a mean-field approach. Whereas the stochastic approach is suitable to a very small timescale, on which stochastic fluctuations in traction forces are observed, the mean-field model addresses a much coarser timescale. On rigid substrates, cycles of loading/unloading occur at a very high frequency. Because the frequency of load-and-fail dynamics is so high, the load-and-fail cycling time is smaller than the minimum timescale addressed by the mean-field model. As a result, the mean-field framework is not able to capture these periodic events on rigid substrates, and its temporal solution reaches steady-state after the first loading event. This allows us to determine the time-averaged traction force 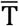, mean number of bound clutches 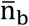 and mean clutch strain rate 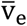 in a more theoretical way by seeking the steady-state solution of Eq. (1). After some algebraic manipulation (see Supplemental Information), we find that the time-averaged probability density of clutch extensions has a double exponential functional form with a very fast decay at a clutch length equal to the clutch rupture length 𝓁_rupt_ = F_b_⁄κ_c_:

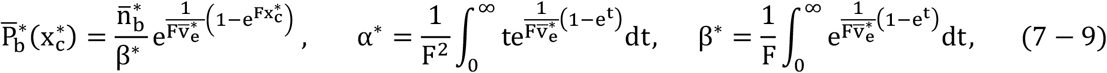

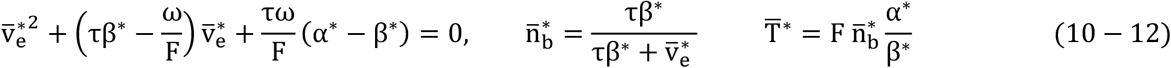

We solve for 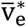 by numerically computing the solution of the nonlinear equation (10). Once we solve for the mean clutch extension rate, we can compute 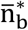 and 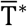 using Eqs. (11) and (12), respectively. We find very good agreement between our mean-field solutions and the solution from the stochastic model, as shown in Fig. (3). We find that traction forces on rigid environments exhibit a biphasic dependence on motor activity (Fig. (3, left)), where the maximum time-averaged traction force is reached at an intermediate value of the myosin activity parameter F^opt^, that monotonically decreases with τ and ω (Fig. (4A)). The time-averaged clutch extension rate also exhibits a maximum at a value of F that is greater than F^opt^ (Fig. (3, right)). Below we derive analytical expressions for 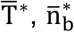 and 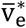 for two asymptotic cases: low-motor activity (F → 0) and high-motor activity (F → ∞).

**FIGURE 3.**
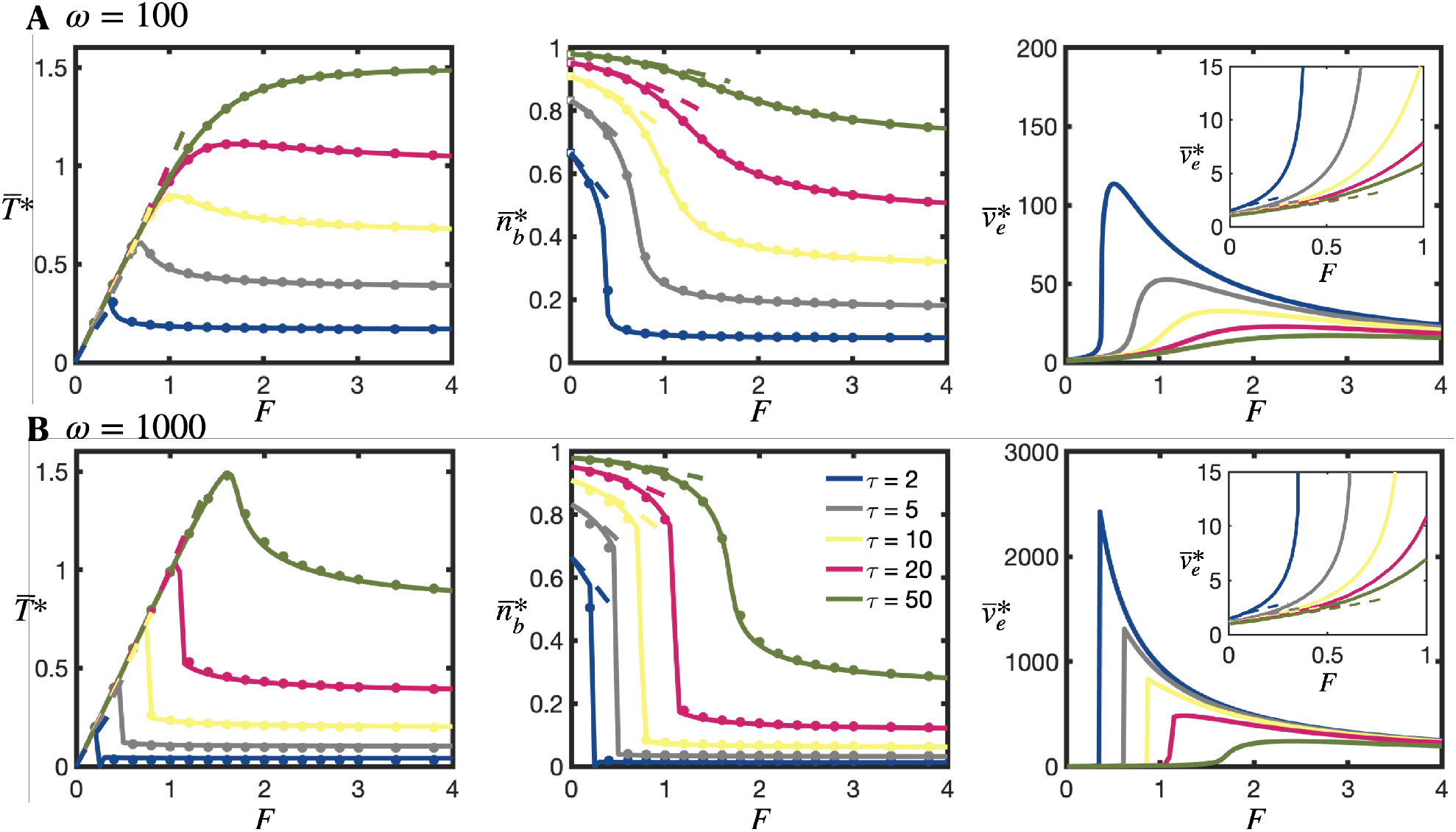
Force transmission on rigid substrates (K → ∞) exhibits a biphasic dependence on motor activity — Time-averaged dimensionless traction force, fraction of bound clutches, and clutch strain rate as a function of the myosin activity parameter F for five different values of the clutch kinetic parameter τ, and two different values of the parameter ω ((A) First row: ω = 100, (B) Second row: ω = 1000). Traction force and fraction of bound clutches monotonically increase with τ, whereas clutch strain rate decreases with τ. Traction force exhibits a maximum at an intermediate value of the myosin activity parameter F^opt^ (see Fig. 4A for further analysis). Clutch extension rate also exhibits a maximum at an intermediate value of the myosin activity parameter F > F^opt^. Curves reach F-independent asymptotic values at high myosin activity (large F) (see Figs. (4B) and (4C) for more details). Solid lines correspond to the numerical solution of the mean-field model (Eqs. (7–12)) and symbols correspond to the numerical solution of the stochastic model. Dashed lines correspond to our low-F asymptotic solutions (Eqs. (13), (S48) and (S49)).

**FIGURE 4.**
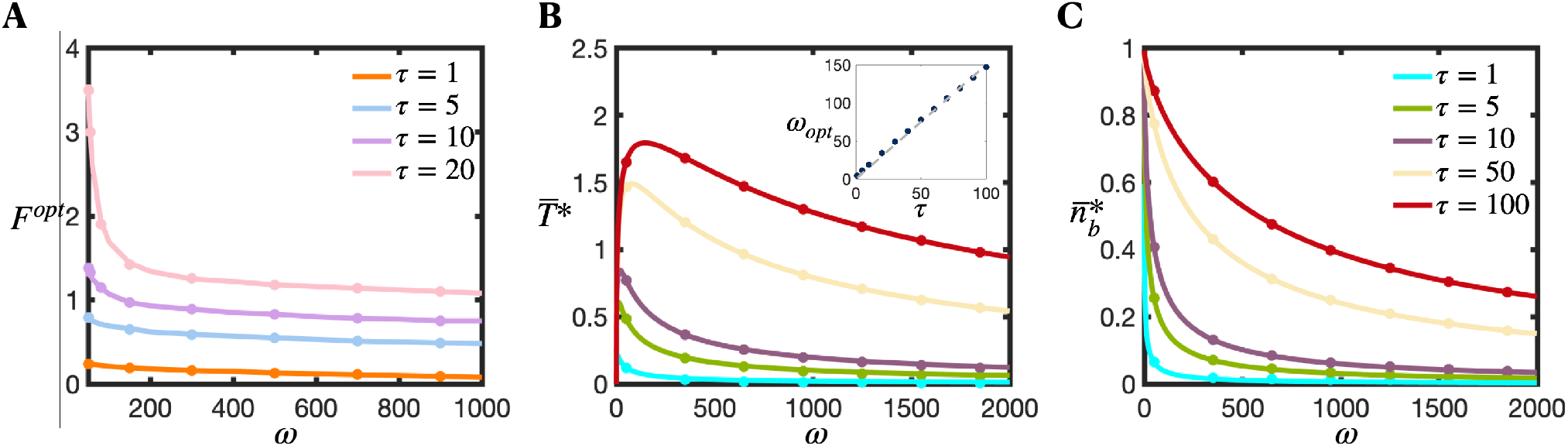
Traction force produced by a strong-motor protrusion on rigid substrates reaches a maximum at an intermediate value of the myosin load-free velocity parameter ω — (A) Optimum myosin activity parameter for maximum traction force on rigid substrates as a function of ω for four different values of the clutch kinetic parameter τ. The optimum activity F^opt^ monotonically decreases with τ and ω. (B, C) Dimensionless time-averaged traction force (B) and average fraction of bound clutches (C) for a strong-motor protrusion (F → ∞) on rigid substrates (K → ∞), as a function of ω for five different values of τ. Force transmission shows a biphasic dependence on the parameter ω, with an optimum ω^opt^ that increases linearly with τ (see inset in (B)). Number of bound clutches increases monotonically with τ and ω. Solid lines correspond to the analytical solutions of the mean-field model (Eqs. (15) and (16)), and symbols are the solutions of the stochastic model. Perfect agreement is found between analytical and stochastic solutions.

#### Low-motor activity (F → 0)

We derived analytical expressions for the time-averaged clutch elongation rate, number of bound clutches and traction force by using perturbation theory (see Supplemental Information). The traction force produced by the protrusion reads:

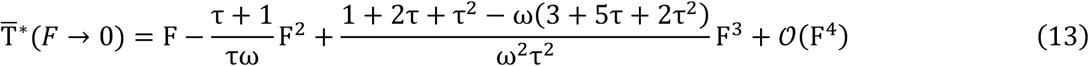

The analytical expressions for the time-averaged clutch elongation rate and number of bound clutches are included in Eqs. (S46) and (S49), respectively. The derived analytical expressions agree very well with numerical solutions, as shown in Fig. (3). The leading order traction force term in Eq. (13) corresponds to the total myosin stall force — that is, in the limit of low myosin activity the time-averaged traction force produced by the protrusion is equal to the maximum available force, as previously explained. The second leading order term in Eq. (13) indicates that as the number of motors increase, the traction force will negatively deviate from the total myosin stall force. An increase in the number of clutches n_c_, the clutch stiffness κ_c_, the clutch association rate constant, k_on_, and the myosin load-free velocity, v_u_, increases the production of traction forces, whereas an increase in the unloaded clutch dissociation rate constant, k_off_, negatively contributes to traction force production. Among all the parameters, the least obvious dependence is that of traction force on myosin load-free velocity. On rigid substrates, actomyosin pulling forces mainly deform molecular clutches, as rigid substrates barely undergo any deformation. In low-motor protrusions, adhesions build strong clusters that inhibit retrograde flows that result in lower clutch loading rates. Consequently, clutches dissociate stochastically before reaching their rupture length giving rise to poor force transmission. A higher value of v_u_ allows clutches to work at their fullest capacity, resulting in stronger retrograde flows, thus higher time-averaged traction forces.

#### High-motor activity (F → ∞)

We also obtained analytical expressions for 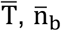 and 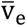 for motor-dominated protrusions. Using perturbation theory, we determine the leading-order terms (see Supplemental Information):

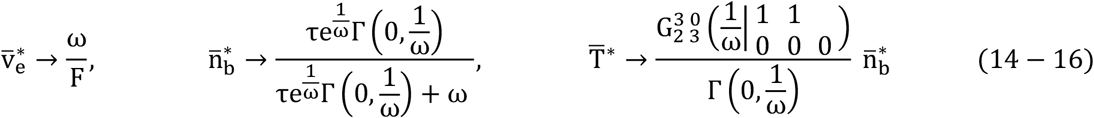

The derived analytical expressions agree very well with our numerical solutions, as shown in Fig. (4B) and (4C). Equation (16) implies that clutches of motor-dominated protrusions elongate on rigid substrates at an average rate that approaches the myosin load-free velocity 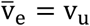. The derived expression for 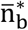 indicates that the fraction of bound clutches monotonically increases as the clutch kinetic parameter τ increases and/or the parameter ω decreases. The derived expression for 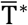 is plotted in Fig. (4B) as a function of ω for different values of τ, and perfect agreement is found between the derived solution and the stochastic model solution. Interestingly, the 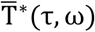 curves reach a maximum value at an optimum value ω^opt^. This optimum dimensionless load-free velocity exhibits a nearly linear dependence with τ, as shown in the inset of Fig. (4B). Therefore, an intermediate value of κ_c_v_u_/F_b_k_off_ exists that maximizes traction forces, and scales linearly with the ratio k_on_/k_off_.

### Clutch reinforcement shifts the optimum substrate stiffness and optimum clutch stiffness to stiffer substrates and stiffer clutches

So far, we have assumed that clutches behave as slip bonds. In some systems (36,37), clutch connections behave as catch bonds with reinforcement instead. In this section, we explore the effect of clutch adhesion reinforcement on force transmission (14,37-41). We assume that the effective clutch association rate linearly depends on the actual fraction of clutches that are bound and whose load exceeds the clutch reinforcement threshold force F_th_ (38). The conservation equation for the probability density thus reads:

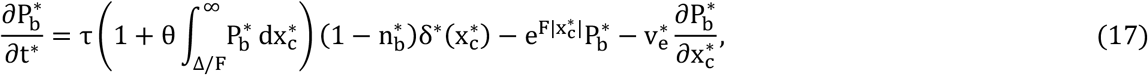

where we have introduced a clutch reinforcement parameter θ and dimensionless clutch reinforcement threshold force ∆= F_th_/F_b_. The dimensionless clutch strain rate 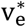 has been defined in Eq. (4). We numerically solve Eq. (17) and find that, in protrusions with a small unloaded myosin velocity parameter ω and low number of myosin motors over clutches, i.e. low F, clutch reinforcement prevents frictional slippage on rigid substrates and force transmission increases monotonically with substrate stiffness, as shown in Fig. (5A). Motor-dominated protrusions, i.e. high F, are associated with large clutch force loading rates due to large myosin pulling forces, and would require an anomalously high clutch reinforcement parameter to strengthen clutches, prevent frictional slippage and allow the production of substantial traction forces on rigid substrates, as manifested by the low traction force transmitted on rigid substrates (Fig. 5B). For larger values of the unloaded myosin velocity parameter, the traction vs substrate stiffness curves are monotonically increasing for sufficiently high values of the clutch reinforcement parameter θ, whereas they display a biphasic behavior for low values of θ, as shown in Fig. (S5A). We find that force transmission on rigid substrates monotonically decrease with an increase in the characteristic clutch reinforcement force ∆, and it can be strongly influenced by the value of ∆, particularly for sufficiently high θ values and low enough ω values, as depicted in Figs. (5C) and (S5B). Clutch reinforcement shifts the optimum substrate stiffness for maximal force production to stiffer substrates, as shown by Figs. (5D) and (S5D), and this optimum stiffness shift is significant only for low values of the unloaded myosin velocity parameter ω, as Fig. (S5C) shows. We further explore analytically the dependence of force transmission on clutch reinforcement for protrusions on rigid substrates. To simplify the analysis, we assume that clutch reinforcement kicks in at vanishing force (∆= 0). After some algebraic manipulation, we find that the dimensionless mean clutch strain rate satisfies the following equation:

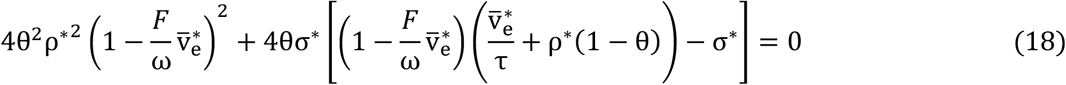

**FIGURE 5.**
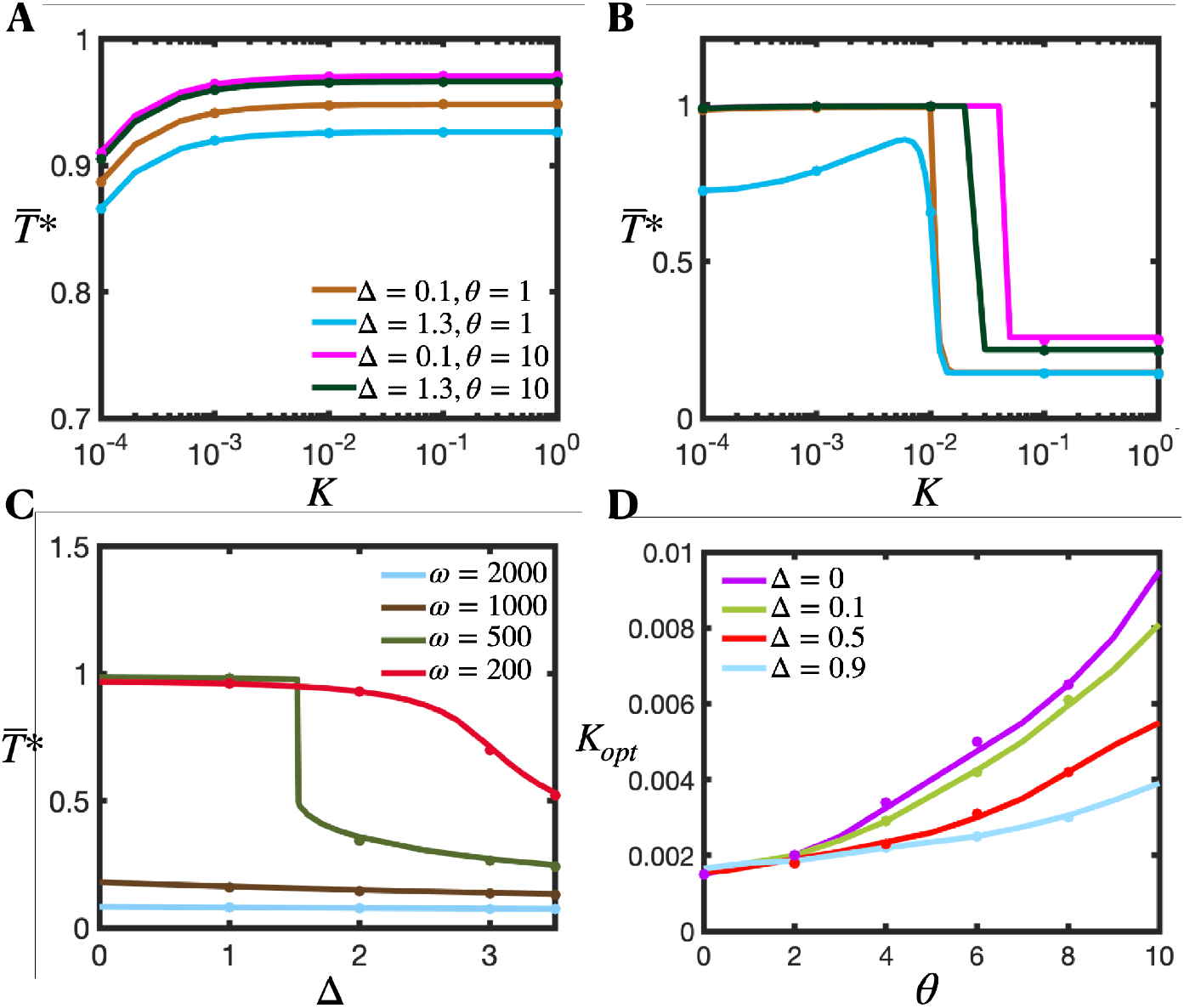
Force transmission on rigid substrates results from a competition between clutch reinforcement and myosin-mediated clutch loading rates. (A-B) Dimensionless time-averaged traction force as a function of substrate stiffness for different values of the clutch reinforcement parameter θ and clutch reinforcement threshold force Δ. Parameter values: *τ* = 1, ω = 200, (A) *F* = 1 and (B) *F* = 10. (C) Dimensionless time-averaged traction force on rigid substrates (K → ∞) as a function of Δ for four different values of the unloaded velocity parameter ω. Parameter values: θ = 10, *F* = 1, *τ* = 5. (D) Optimum substrate stiffness as a function of θ for different values of Δ. Parameter values: ω = 200, *F* = 1, *τ* = 1. Solid lines are the numerical solution of the developed mean-field model (Eq. 17), and symbols are the numerical solutions of the stochastic motor-clutch model.

The dimensionless time-averaged number of bound clutches and traction force read:

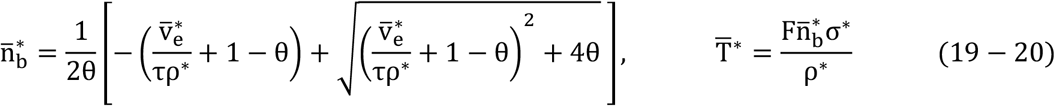

where

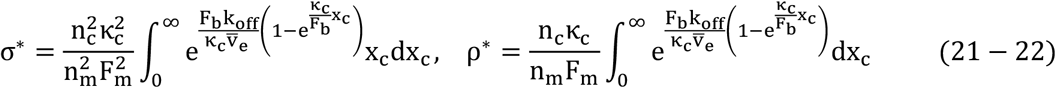

We solve Eq. (18) numerically and find that the mean traction force produced by a single protrusion on rigid substrates is significantly enhanced by clutch reinforcement for protrusions with fast clutch association kinetics (large τ), as shown in Fig. (6A). As the clutch reinforcement parameter θ increases, the time-averaged traction force produced by the protrusion rises, and it eventually reaches a plateau at large values of θ, where the protrusion nearly reaches stall conditions, as shown in Fig. (6A) for τ = 3 and τ = 5. We showed in Fig. (4B) that, in the absence of clutch reinforcement, the traction force curves reach a maximum value at an optimum value ω^opt^. Our mean-field model with clutch reinforcement shows that adhesion reinforcement shifts the optimum unloaded velocity parameter to larger values as depicted in Fig. (6B). We also find that clutch reinforcement on rigid substrates shifts the optimum clutch stiffness to higher stiffnesses, as Fig. (6C) shows. Interestingly, the shifts in the optimum unloaded velocity parameter and clutch stiffness are very sensitive to changes in the clutch reinforcement parameter θ for the largest values of the clutch kinetic parameter τ explored, whereas they are nearly independent of θ for the smallest value of τ explored (τ = 1), as Figs. (6B) and (6C) show, respectively. The sensitivity of force transmission on the parameter θ for different myosin activity levels is quantified in Fig. (6D). We find that low-motor activity protrusions (small F) produce θ-independent traction forces, since the protrusion is already operating at stall conditions. High-motor activity protrusions (large F) display high-frequency load-and-fail dynamics on rigid substrates, and force transmission and mean number of bound clutches are enhanced as the clutch reinforcement parameter increases, as shown in Figs. (6D), (6E) and (6F). The asymptotic leading order term solutions for the time-averaged traction force and mean number of bound clutches for motor-dominated protrusions with reinforcement are obtained by taking the limit F → ∞ on Eqs. (18-22). Our analytical solutions (Eqs. S70-S72) are in very good agreement with the stochastic motor-clutch model solutions, as shown in Figs. (6E) and (6F). We find that high-motor protrusions on rigid substrates can avoid frictional slippage and produce large traction forces provided that the clutch parameters θ and/or τ are sufficiently large and the parameter ω is sufficiently small, as shown in Figs. (6E) and (S6). More details are included in the Supplemental Information. Finally, we explore the role of clutch catch bond behavior on force transmission and find nearly independent traction force generation on catch properties (See Fig. (S7)).

**FIGURE 6.**
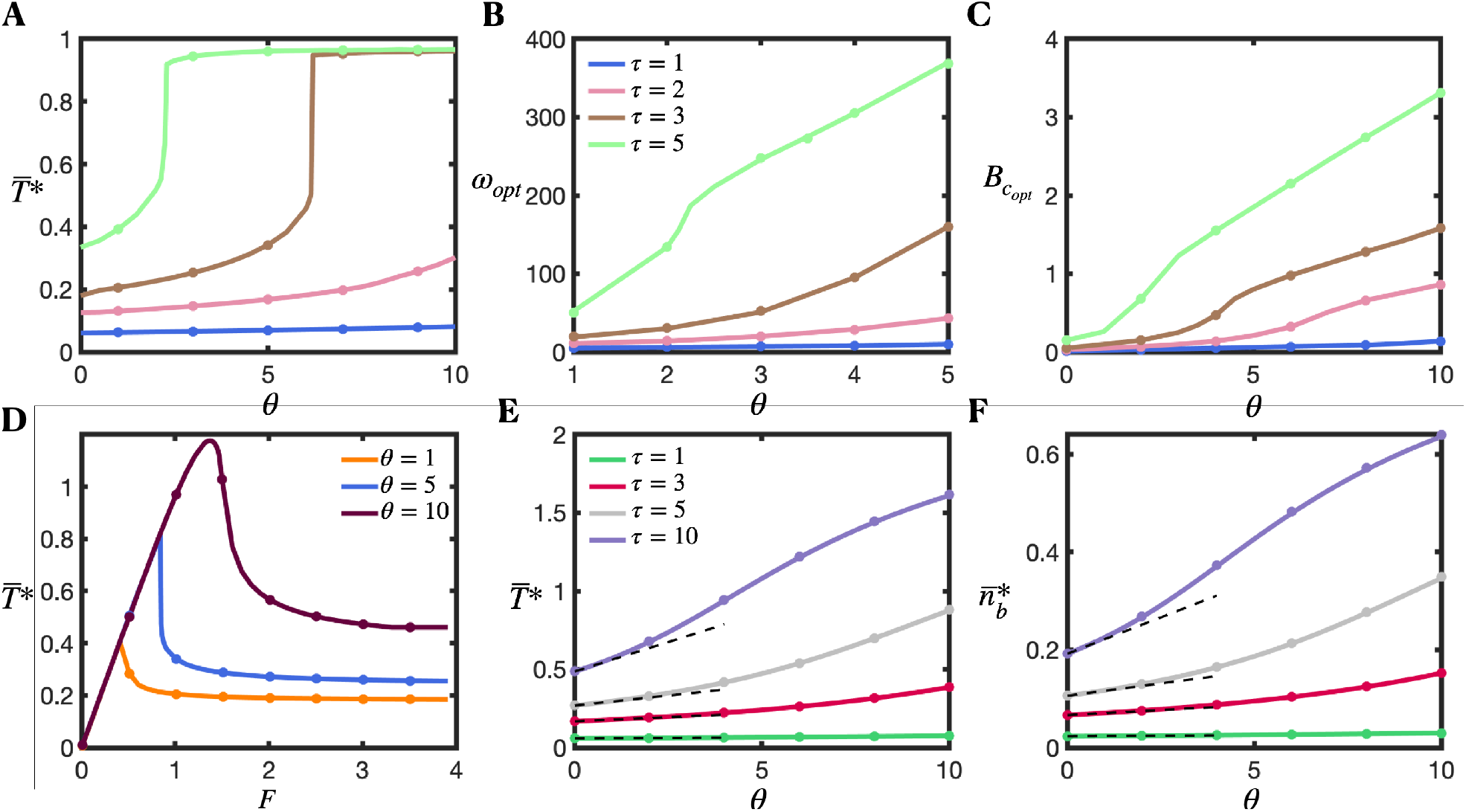
Clutch reinforcement on rigid substrates shifts the optimum unloaded velocity parameter and optimum clutch stiffness to larger values. (A) Time-averaged traction force (parameter values: F = 1, ω = 200, K → ∞), (B) optimum unloaded velocity parameter (parameter values: F = 1, K → ∞), and (C) optimum clutch stiffness parameter (parameter values: F = 1, ω^M^ = 200, K′ = 1) as a function of the clutch reinforcement parameter θ for four different values of the clutch kinetic parameter *τ*. Notice that K = K′/β_c_ and ω = ω′β_c_. Solid lines are the numerical solution of the developed mean-field model (Eq. 18) and symbols are the numerical solutions of the stochastic motor-clutch model. (D) Time-averaged traction force as a function the myosin activity parameter *F* for three different values of the clutch reinforcement parameter. Parameter values: τ = 3, ω = 200, K → ∞. Solid lines are the numerical solution of the mean-field model (Eq. 18) and symbols are the numerical solutions of the stochastic motor-clutch model. (E) Time-averaged traction force and (F) number of bound clutches as a function of the clutch reinforcement parameter for four different values of the clutch kinetic parameter. Parameter values: F → ∞, K → ∞, ω = 200. Solid lines correspond to our analytical solutions (Eqs. (S70) and (S71)), symbols correspond to the numerical solutions of the stochastic motor-clutch model, and black dashed lines are the asymptotic solutions for small θ obtained in Eqs. (S71) and (S73). Δ=0 in all panels.

## DISCUSSION

In the present study, we developed a mean-field model to understand traction force production of adhesion-based cellular protrusions on compliant substrates. We considered cell protrusions to respond to extracellular matrix rigidity by employing a molecular clutch mechanism (2,13), which allows cells to transmit forces to their surrounding matrix. We applied scaling analysis to our model with the purpose of deriving an analytical expression for the optimum substrate stiffness, i.e., the substrate stiffness that maximizes cellular traction forces. We found that maximum traction forces occur when the characteristic time for adhesion molecules to mechanically link actin filaments to the extracellular matrix scale with the characteristic time for adhesion molecules to rupture by force. For substrate stiffnesses higher than the optimal, clutch rupture times become smaller than clutch binding times after a small transient at the beginning of each load-and-fail cycle, causing a clutch failure cascade that terminates the migration cycle, and the protrusion undergoes frictional slippage in a frequent manner (2,16) decreasing the overall force transmission. For substrate stiffnesses lower than the optimal, force transmission loading is slow, and clutches disengage before high force transmission is achieved. We numerically solved the mean-field model equations and found very good agreement between our analytical results, our numerical solutions obtained from the mean-field model and the stochastic model results (2).

We additionally provided a physical explanation for why optimum substrate stiffness is found to be independent of clutch stiffness, as previously reported (10,16), since clutch binding times and clutch rupture times are independent of clutch stiffness near the optimum. Even though the optimum substrate stiffness is not affected by clutch stiffness, we found that traction force generation is strongly influenced by clutch stiffness, especially on rigid substrates, and discovered the existence of an intermediate clutch stiffness for maximum force transmission, a result that could be important for understanding cell behavior on stiff materials such as tissue culture plastic or glass, implanted biomedical devices, and bone. We additionally found that motor-dominated protrusions shift the clutch stiffness optimum towards softer clutch rigidities whereas clutch-dominated protrusions shift it towards stiffer clutch rigidities. In addition, we have found that protrusions on rigid substrates display a biphasic dependence of force transmission on motor activity. This is particularly interesting, since motor activity levels can be modulated by the internal cellular state as well as by environmental cellular conditions. As an example, nuclear deformation mediated by enhanced cellular confinement can activate mechanotransduction pathways (42,43) that enhance actomyosin contractility and therefore control force transmission. Our model predicts that a significant increase in motor activity can dramatically diminish adhesion-based traction forces and therefore cells might require alternative mechanisms to efficiently migrate within highly confined spaces or within other mechanochemical environments that promote high motor activity levels. Our mean-field model also predicts that traction force produced by a strong-motor protrusion on rigid substrates reaches a maximum at an intermediate value of the myosin load-free velocity parameter. This novel result could have important implications in understanding the contribution of different myosin motor types on force transmission, as different myosin motors produce distinct force-velocity curves and have different myosin load-free velocities (44–46). It would also be interesting to test this prediction experimentally, and it encourages the design of motor-oriented therapeutic strategies to modulate traction force production, and potentially cell migration of cells that operate in a high-motor activity cellular state.

We have also shown that strong-motor protrusions produce maximum traction forces at optimum substrate stiffnesses that are myosin-insensitive, that a motor-clutch balanced protrusion is characterized by an optimum substrate stiffness sensitive to myosin activity (as shown in (10)), and that clutch-dominated stalled protrusions are characterized by substrate stiffness-insensitive traction forces as previously shown (10). We explored these three different cellular regimes in depth, derived analytical expressions for the necessary and sufficient conditions required for the cell to operate at each regime, and obtained analytical expressions for the traction force generated by cellular protrusions on non-compliant substrates, in the limits of low and high myosin activity. We found that it is possible to enforce protrusions to operate in the motor-dominated regime when the ratio between the total myosin stall force in the protrusion and the effective maximum clutch elastic force (F) is much higher than the clutch kinetic parameter τ. We also found that the optimum substrate stiffness for cell protrusions that lie in the motor-dominated regime depends proportionally to the number of clutches, clutch association kinetic constant and clutch rupture force and inversely proportional to the unloaded myosin velocity. Our results suggest potential strategies to modulate traction force produced by cells that lie in the motor regime. Our model also predicts that clutch reinforcement can substantially increase the traction force produce by protrusions that display load-and-fail dynamics, and significantly shift the optimum substrate and clutch stiffnesses to larger values. Whereas traction force-substrate stiffness curves were found to be monotonically increasing functions in reinforced protrusions with small unloaded myosin velocities and relatively low number of myosin motors over clutches, motor-dominated protrusions displayed a biphasic dependence of force transmission on substrate stiffness for the clutch reinforcement strengths explored, requiring anomalously high clutch reinforcement strengths to prevent frictional slippage and to allow the production of significant traction forces on rigid substrates. These results provide key insights to effectively design novel cell and stromal engineering strategies to strengthen or weaken clutch reinforcement with the aim of, respectively, enhancing or decreasing traction force production and potentially cell migration capabilities. Finally, we showed that protrusions with clutch catch bond properties can modulate force transmission but do not alter the optimum substrate stiffness for maximum force.

Our current study has some limitations. We have assumed that the mechanical properties of clutches are those of the weakest link in the adhesion complex, a standard assumption in the motor-clutch framework. The constitutive adhesion proteins that form clutch complexes flow retrogradely at different speeds (47,48), with actin-binding proteins flowing at high speeds and more coherently, matrix-binding proteins flowing at low speeds and mostly incoherently, and core clutch proteins linking actin-binding and matrix-binding proteins flowing at intermediate speeds and with somewhat coherent motion. The reported differences in retrograde flow speeds agree well with the motor-clutch hypothesis, and this could imply the absence of a clear “weakest link”, and that the constitutive protein-protein adhesion bonds may have similar mechanical properties with multiple bonds influencing force transmission. The reported differential coherence could suggest that adhesion complex links undergo slippage and do not maintain robust connections (49), which could limit force transmission and efficiency. We have also assumed that all bound clutches equally deform at any instant in time, a reasonable assumption in the presence of high density of actin crosslinkers and large amounts of myosin heads and clutches compared with number of actin filaments. In this way, myosin forces will be approximately equally distributed among all actin filaments and the same number of clutches will approximately bind to each actin filament. Multiple clutch-actin connections as well as the effect of actin crosslinkers are additional features that could be included in future continuum models for traction force production. We have also assumed that the substrate is a Hookean elastic material. However, tissues and extracellular fibril networks are not linearly elastic, but display complex mechanical behaviors, including nonlinear elasticity such as stress stiffening and stress softening (50,51), viscoelasticity (52,53) and mechanical plasticity (54). An experimental and theoretical study showed that soft substrates with faster stress relaxation times enhance cell migration and increase filopodia length and lifetime (55), whereas a motor-clutch framework combined with a simple viscoelastic solid model for the substrate reported that maximum single protrusion front speeds are achieved at intermediate relaxation times in soft matrices (34). Force loading rates are very slow in soft substrates and viscous dissipation provides additional resistance to retrograde myosin pulling forces that increases the initial force loading rate, reducing actin retrograde flows, and increasing membrane front speeds and cell migration. Our analysis also assumes that forces are applied at a singular point, whereas in reality, they are spatially distributed, which affects the effective stiffness of the environment (56). The mechanical response of the extracellular space to cellular forces plays a critical role in force transmission and cell migration; and determining the constitutive mechanical model that better describes the extracellular space is a potential opportunity for future experimental and theoretical studies.

Overall, our study provides new insights of force transmission of adhesion-based cellular protrusions on compliant elastic substrates and provides a quantitative analysis of regulation of force transmission by modulation of cellular components and extracellular rigidity. More specifically, we identify the existence of an optimum clutch stiffness for maximum traction force and report the existence of an intermediate myosin mean stall force and unloaded myosin velocity for maximum force transmission on rigid substrates. The results predicted by our model on rigid substrates are particularly interesting due to the difficulties that arise during the quantification of traction force measurements on low-compliant extracellular environments with traditional methods such as traction force microscopy. The reported biphasic dependence of force transmission on motor activity and unloaded motor velocity could be potentially tested via novel molecular tension sensor experiments and opens new research avenues in biological systems with stiff cellular environments. Unraveling the mechanics of cellular force production and migration in stiff cellular environments is crucial in many aspects of health and human physiology. Cell migration dysregulation in stiff environments is associated with many processes in development and human diseases, such as clinical complications, tissue remodeling and immune modulation in inflammatory responses near implanted medical devices (57,58), bone cancer metastasis (59), fibrosis in cancer (60,61), calcified bone remodeling processes and homeostasis (62) and dental lamina development and degradation processes (63), among others. Additional studies are needed to identify the main adhesion clutch constituents, as well as the key mechanisms for efficient cellular force transmission/migration as well as activation of mechanotransduction pathways in different complex mechanochemical microenvironments. The mechanotransduction of cellular forces transmitted to the extracellular fibril network into biochemical signals leads to transcriptional regulation in the nucleus (64–66) that can modulate force transmission, thus influencing cell migration in non-trivial ways. Further understanding will speed up the discovery of new therapeutic strategies to treat human disease for improved patient outcomes.

## AUTHOR CONTRIBUTIONS

R.A.M. designed research, developed the biophysical model, performed mathematical derivations and numerical simulations, and wrote the manuscript. P.P.P and D.O. secured funding and oversaw all aspects of the study, and provided edits to the manuscript.

## DECLARATION OF INTERESTS

Dr. Provenzano is a member of the Scientific Advisory Board for Parthenon Therapeutics. The other authors declare no competing interests.

## ACKNOWLEDGEMENTS

Research reported in this publication was supported by grants U54 - CA210190, P01 - CA254849, and U54 - CA268069. We thank members of the Provenzano and Odde labs for helpful conversations throughout the course of this work. The content of this work is solely the responsibility of the authors and does not necessarily represent the official views of the NIH.

---

**FIGURE S1.**
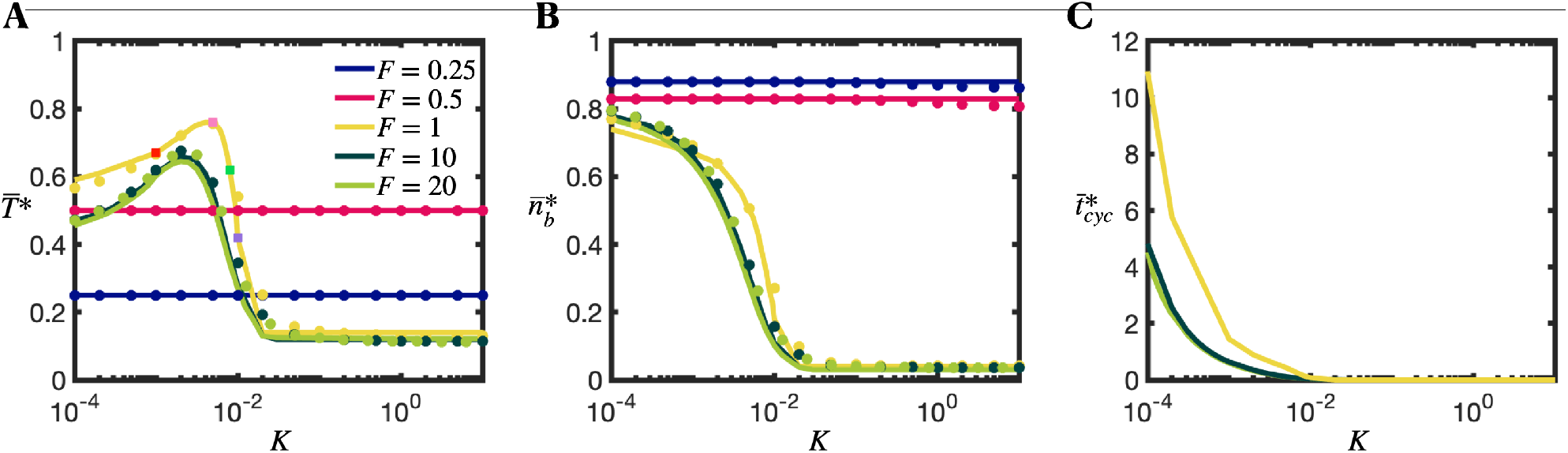
Traction force production of individual cellular protrusions exhibit three different regimes: a motor-dominated regime, an intermediate regime, and a clutch-dominated regime. Force transmission is sensitive to substrate compliance — (A) Dimensionless time-averaged traction force 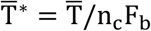, (B) time-averaged fraction of bound clutches 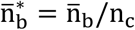, and (C) dimensionless time-averaged cycling time 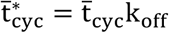 as a function of the dimensionless substrate stiffness K for various values of the myosin activity parameter F. Protrusions can belong to three different regimes: a motor-dominated regime characterized by the existence of an optimum stiffness for maximal traction that is largely independent of myosin activity (F = 10 and F = 20 in part A), a clutch-dominated regime characterized by stiffness-independent traction forces (F = 0.25 and F = 0.5 in part A), and an intermediate regime characterized by an optimum stiffness sensitive to motor activity (F∼1 in part A). The three regimes are analyzed in detail in the main text. Solid lines are the numerical solution of the mean-field model equations (3) and (4), and circular solid symbols are the mean statistics obtained from the numerical solution of the stochastic model. There is a very good agreement between our mean-field model solution and stochastic model solution. Parameter values: τ = 10, ω = 2000.

**FIGURE S2.**
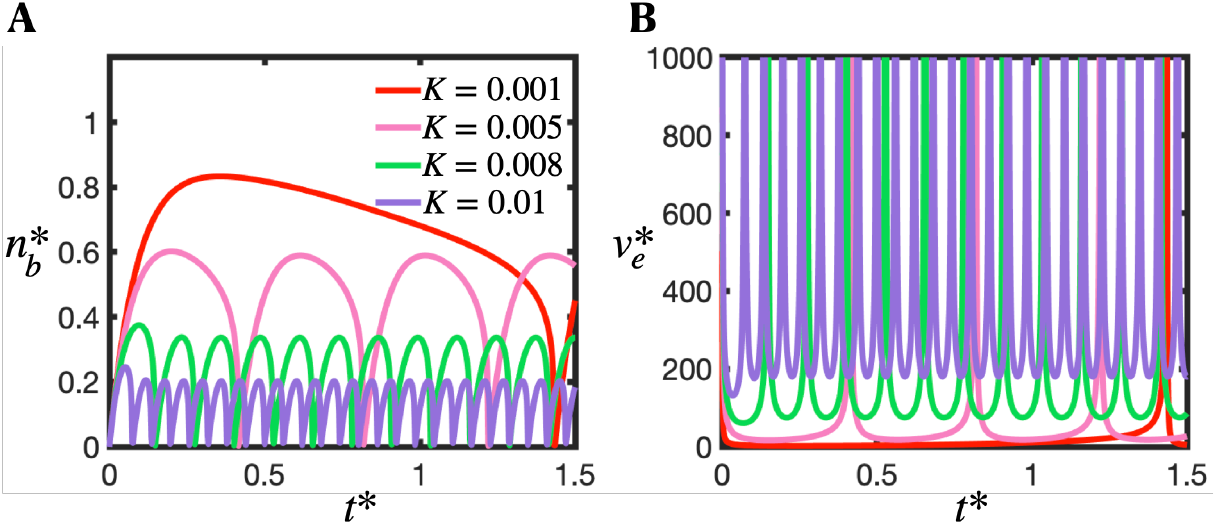
Protrusions in the motor-clutch balanced regime display load-and-fail dynamics. Time-evolution of fraction of bound clutches (A) and clutch extension rate (B) obtained by our mean-field model for 4 different substrate stiffnesses. Parameter values: F = 1, τ = 10, ω = 2000.

**FIGURE S3.**
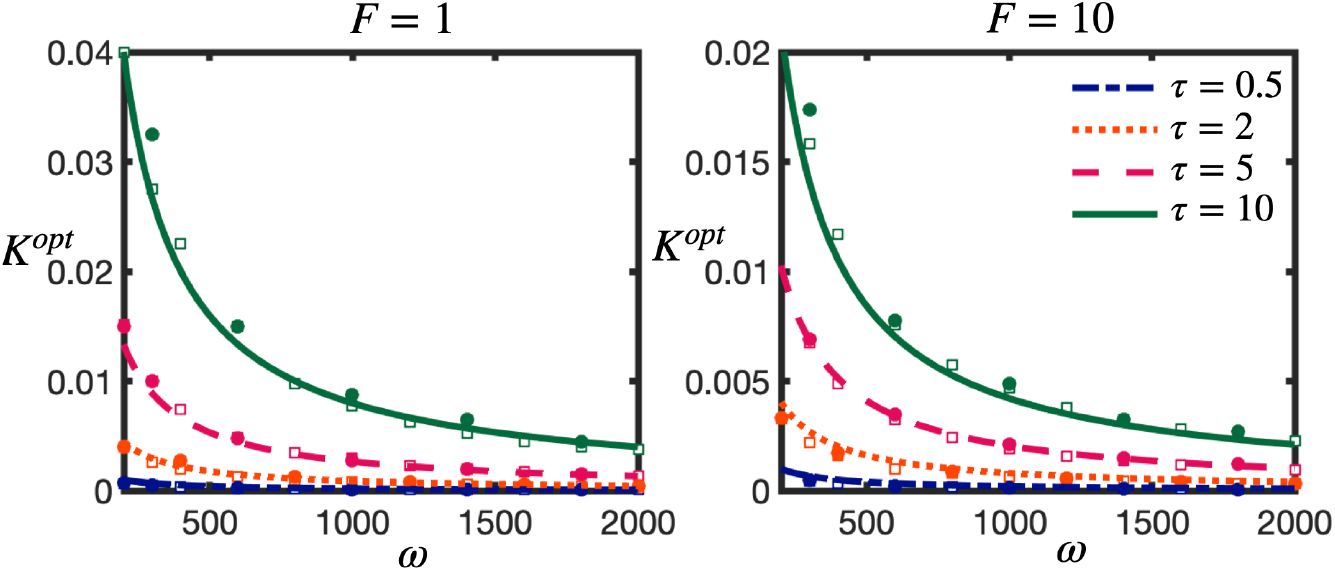
Dimensionless optimum substrate stiffness 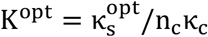 as a function of the dimensionless myosin load-free velocity ω, for two different values of the myosin activity parameter F. Solid lines correspond to our derived analytical solution (Eq. 6), open symbols correspond to the numerical solution of the mean-field model (Eqs. (3) and (4)), and closed symbols correspond to the numerical solution of the stochastic model.

**FIGURE S4.**
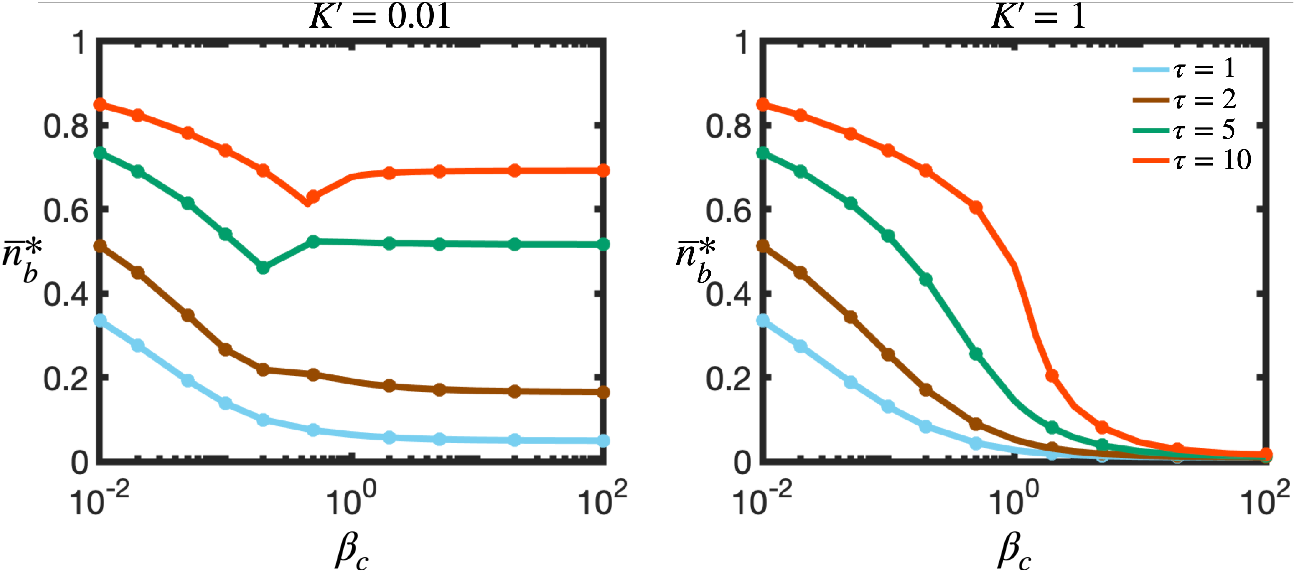
Dimensionless time-averaged number of bound clutches as a function of the clutch stiffness parameter β_c_ for different values of the clutch kinetic parameter τ and for two values of the substrate stiffness parameter *K*′ (*K* = *K*′/β_c_): (left) *K*′ = 0.01 and (right) *K*′ = 1. Parameter values: *F* = 1, *ω*′ = 200 (*ω* = *ω*′β_c_). Notice that κ_c_ ∝ β_c_.

**FIGURE S5.**
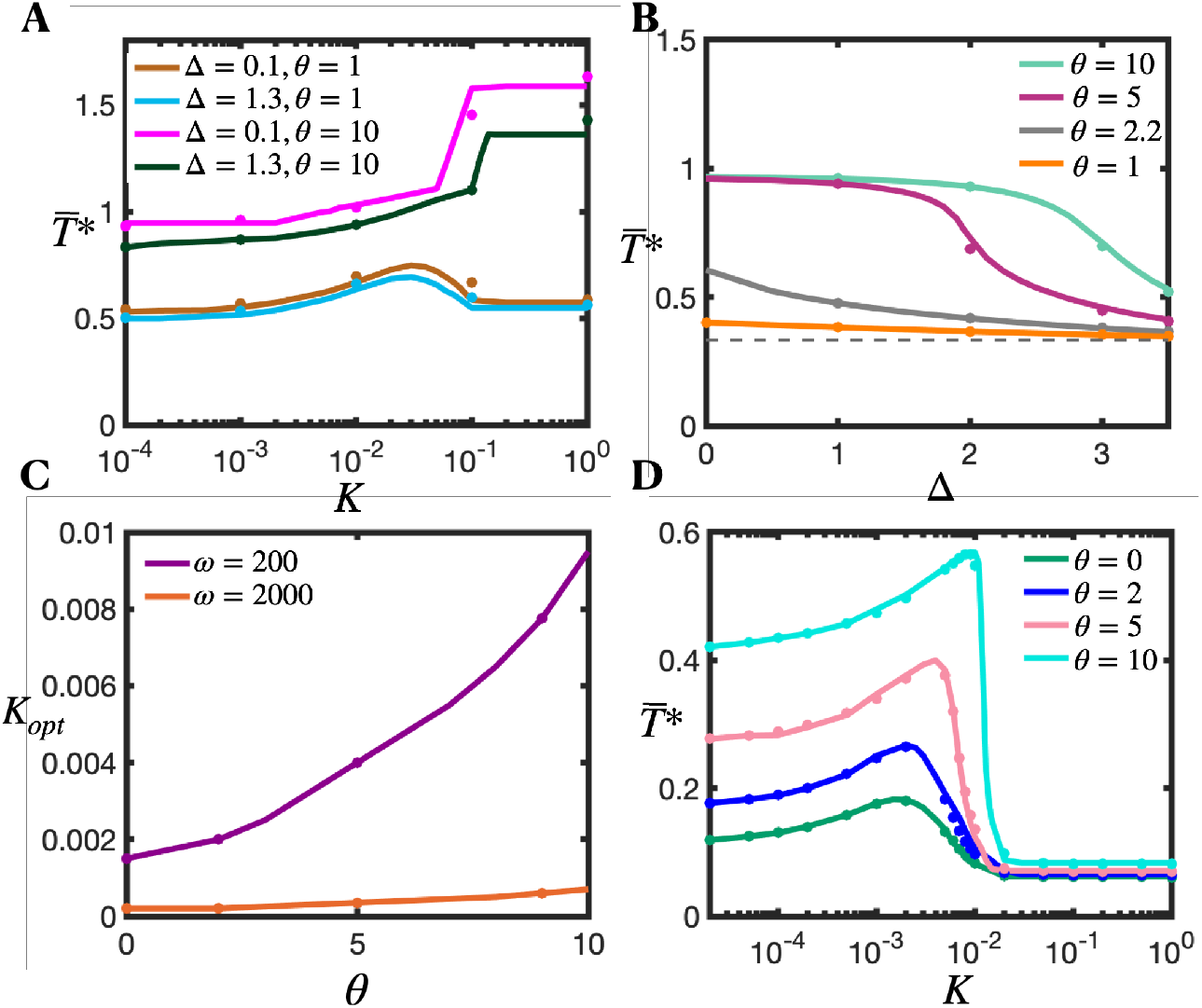
Clutch reinforcement shifts the optimum substrate stiffness for maximal force transmission to stiffer substrates. (A) Dimensionless time-averaged traction force as a function of substrate stiffness for two values of the clutch reinforcement parameter θ and clutch reinforcement threshold force Δ. Parameter values: *τ* = 1, ω = 200, *F* = 10. (B) Dimensionless time-averaged traction force on rigid substrates (K → ∞) as a function of the dimensionless clutch reinforcement threshold force Δ for four different values of the clutch reinforcement parameter. Dashed gray line corresponds to the force transmission curve in the absence of clutch reinforcement (θ = 0). Parameter values: ω = 200, *F* = 10, *τ* = 5. (C) Optimum substrate stiffness as a function of the clutch reinforcement parameter for two different values of the dimensionless myosin unloaded velocity ω. Solid lines are the numerical solution of the developed mean-field model (Eq. 18) and symbols are the numerical solutions of the stochastic motor-clutch model. Parameter values: ω = 200, *F* = 1, *τ* = 1. (D) Dimensionless time-averaged traction force as a function of the dimensionless substrate stiffness for four different values of the clutch reinforcement parameter θ. Parameter values: ω = 200, *F* = 1, *τ* = 1, Δ= 0.

**FIGURE S6.**
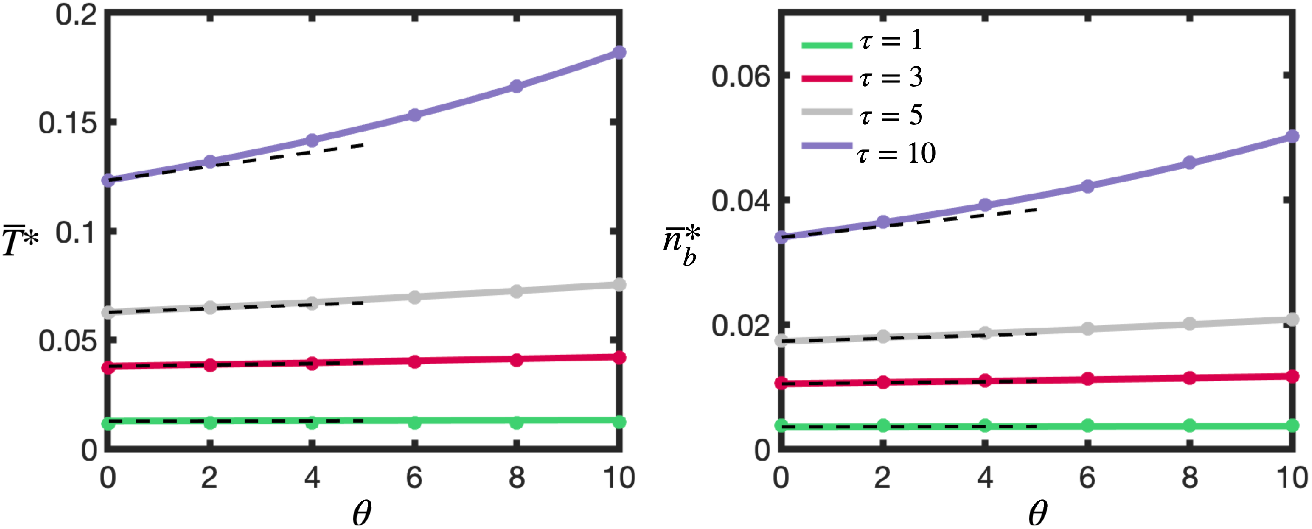
Time-averaged traction force (left) and number of bound clutches (right) as a function of the clutch reinforcement parameter θ for four different values of the clutch kinetic parameter τ. Parameter values: F → ∞, K → ∞, ω = 2000, ∆= 0. Solid lines correspond to the analytical solution (Eqs. (S70) and (S71)), symbols correspond to the numerical solutions of the stochastic motor-clutch model and black dashed lines are the asymptotic solutions when θ → 0 obtained in Eqs. (S71) and (S73).

**FIGURE S7.**
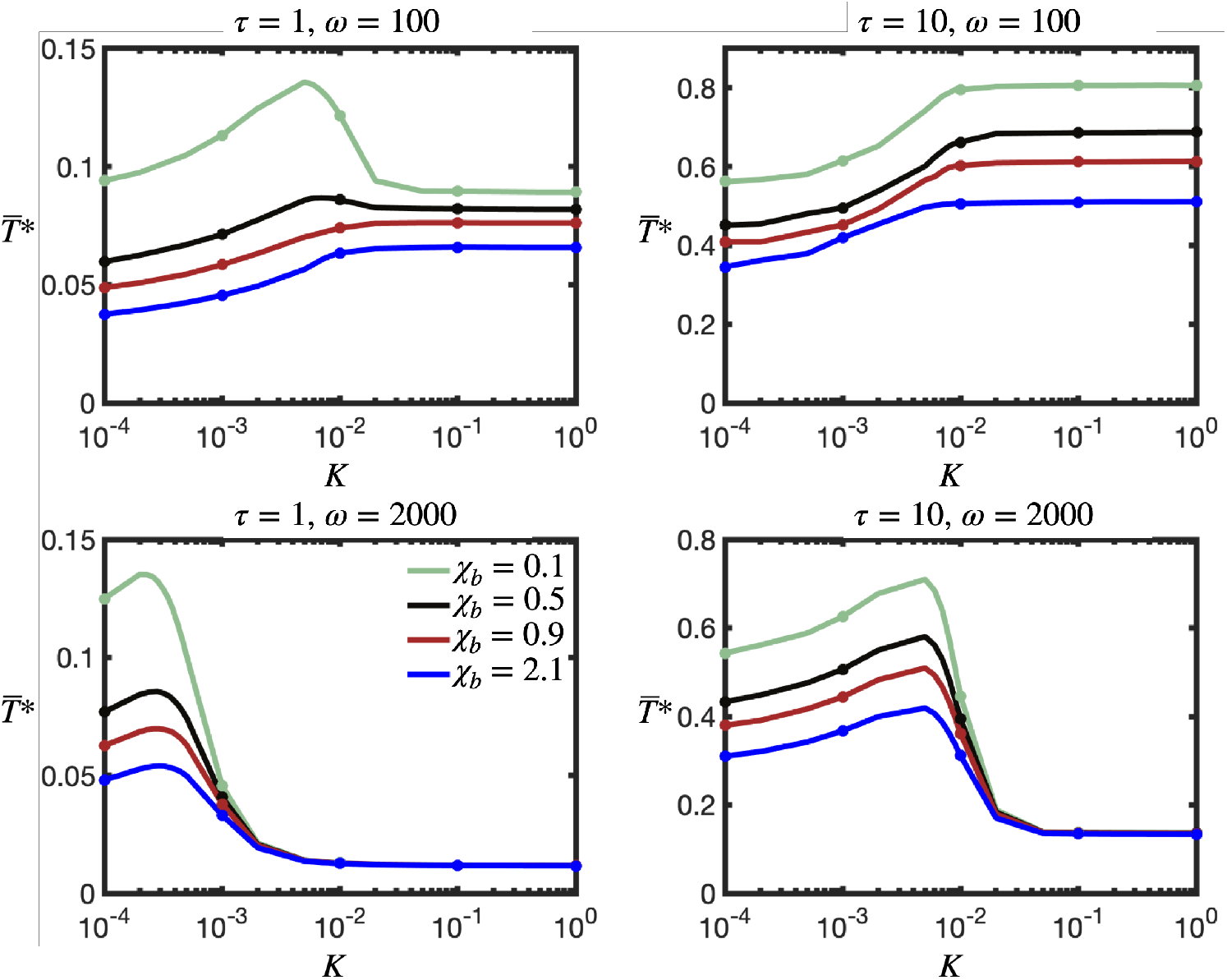
Optimum substrate stiffness for maximal for transmission is independent of the catch bond force parameter χ_b_. Time-averaged traction force as a function of substrate stiffness for four different values of χ_b_. Parameter values: F = 1, K_off_ = 20. Solid lines correspond to the mean-field numerical solution (Eq. (S87)) and symbols correspond to the numerical solutions of the stochastic motor-clutch model

