## Supplemental Information for "“Optimality of motor and clutch mechanical properties in a generalized model for cell traction forces”"

**SUPLEMENTAL INFORMATION**

**Mean-field conservation equation of the probability density**

The Markovian nature of clutch dynamics allow us to relate the probability density $P_b$ at time $t + \Delta t$ to its value at an earlier time $t$ with an integral equation of the form

$$P_b(x_c, t + \Delta t) = k_{on}(n_c - n_b)\delta(x_c)\Delta t - k_{off}^{load} P_b(x_c, t)\Delta t$$
$$+ \int_{-\infty}^{\infty} d(\Delta x_c)\, P_b(x_c - \Delta x_c, t)\phi(x_c - \Delta x_c|\Delta x_c, \Delta t). \qquad (S1)$$

The first term on the RHS of Eq. (S1) corresponds to clutch binding kinetics, where the Dirac delta function $\delta(x_c)$ has been included to specify that the clutch length at the binding time is equal to its resting length. The second term on the RHS of Eq. (S1) accounts for clutch unbinding kinetics, where the clutch lifetime decreases exponentially by force according to Bell's law $k_{off}^{load} = k_{off}\, e^{\kappa_c|x_c|/F_b}$. The third term on the RHS of Eq. (S1) accounts for clutch extensions, where $\phi(x_c|\Delta x_c, \Delta t)$ is the probability of clutches with extension $x_c$ undergoing an extension $\Delta x_c$ during a time $\Delta t$. Ignoring the binding/unbinding kinetic terms, Eq. (S1) says that the probability density of clutches to be with extension $x_c$ at time $t + \Delta t$ is given by the product of the probability density for clutches to be with extension $x_c - \Delta x_c$ at time $t$, multiplied by the probability of experiencing an extension $\Delta x_c$ during a period of time $\Delta t$, and integrated over all possible extensions. Notice that the probability of experiencing a clutch extension of any magnitude is normalized to 1:

$$\int_{-\infty}^{\infty} d(\Delta x_c)\, \phi(x_c - \Delta x_c|\Delta x_c, \Delta t) = 1. \qquad (S2)$$

We expand $P_b(x_c - \Delta x_c, t)$ and $\phi(x_c - \Delta x_c|\Delta x_c, \Delta t)$ in Taylor series about $x_c$, and truncate the series to first order. We get,

$$P_b(x_c, t + \Delta t) = k_{on}(n_c - n_b)\delta(x_c)\Delta t - k_{off}^{*} P_b(x_c, t)\Delta t + P_b(x_c, t) - \frac{\partial}{\partial x_c}(P_b\overline{\Delta x_c}), \qquad (S3)$$

where the mean clutch extension during a time $\Delta t$ is

$$\overline{\Delta x}_c(\Delta t) = \int_{-\infty}^{\infty} d(\Delta x_c)\, \Delta x_c \phi(x_c|\Delta x_c, \Delta t) = v_e \Delta t, \qquad (S4)$$

The clutch deformation rate $v_e$ is equal to the difference between the actin retrograde flow velocity and the substrate deformation rate. Rearranging terms in Eq. (S3) and taking the limit $\Delta t \to 0$ allows us to obtain the conservation equation for the probability density

$$\frac{\partial P_b}{\partial t} = k_{on}(n_c - n_b)\delta(x_c) - k_{off}\, e^{\kappa_c x_c/F_b} P_b(x_c, t) - v_e \frac{\partial P_b}{\partial x_c}. \quad (S5)$$

An equivalent governing equation has been obtained in previous studies (1,2) by using standard mean-field approximations, where they explore steady-state solutions of single protrusions on noncompliant substrates.

**Derivation of the optimum substrate stiffness for maximum traction force**

We proceed to estimate the clutch binding time $t_{bind}$ and clutch rupture time $t_{rupt}$ to obtain an analytical expression for the optimum substrate stiffness that maximizes traction forces. We carry out the mathematical derivation in dimensional form so as not to lose physical intuition throughout the process. We take the zeroth moment of Eq. (1) to obtain the approximated time evolution of the number of bound clutches before clutch bonds break due to load

$$\frac{dn_b}{dt} \approx k_{on} n_c - (k_{on} + k_{off})\, n_b, \qquad t < t_{rupt} \qquad (S6)$$

The number of bound clutches thus scales as $n_b \sim (k_{on}/k_{on} + k_{off}) n_c$. The clutch binding time — that is, the characteristic time at which the number of clutches that link the actin cytoskeleton to the extracellular medium reaches equilibrium, scales as $t_{bind} \sim 1/k_{on} + k_{off}$. We define the clutch rupture time $t_{rupt}$ as the ratio between the characteristic clutch rupture length $\ell_{rupt} = F_b/\kappa_c$ and the characteristic clutch extension rate $v_{e_c}$: $t_{rupt} = \ell_{rupt}/v_{e_c}$. We now proceed to estimate $v_{e_c}$. Before clutches dissociate due to force, the first-order moment and the clutch extension rate approximately satisfy the following relations

$$\frac{d\ell_b}{dt} \approx v_e n_b - k_{off}\ell_b, \qquad t < t_{rupt}, \tag{S7}$$

$$(\kappa_s + n_b\kappa_c)v_e \approx v_u\kappa_s - \left(\frac{v_u\kappa_s}{n_m F_m} - k_{off}\right)\kappa_c\ell_b, \qquad t < t_{rupt} \tag{S8}$$

where we have used Eq. (2). We take the derivative of Eq. (S8) with respect to time, use Eqs. (S6) and (S8), and rearrange terms to get the following ordinary differential equation for the clutch elongation rate, valid for $t < t_{rupt}$,

$$\frac{dv_e}{dt} + \frac{n_b\kappa_c}{(\kappa_s + n_b\kappa_c)}\left[k_{on}\left(\frac{n_c - n_b}{n_b}\right) + \frac{v_u\kappa_s}{n_m F_m} + k_{off}\left(\frac{\kappa_s}{n_b\kappa_c} - 1\right)\right]v_e \approx \frac{n_b\kappa_c\kappa_s k_{off} v_u}{(\kappa_s + n_b\kappa_c)^2}\left(1 - \frac{\kappa_s}{n_b\kappa_c}\right). \tag{S9}$$

After a small transient from the onset of loading and for not very rigid substrates, it is expected that enough clutches will bind so that the substrate rigidity is much smaller than the ensemble clutch stiffness ($\kappa_s \ll n_b\kappa_c$). Under this assumption, Eq. (23) simplifies as

$$\frac{dv_e}{dt} + \left[k_{on}\left(\frac{n_c - n_b}{n_b}\right) + \frac{v_u\kappa_s}{n_m F_m} - k_{off}\right]v_e \approx \frac{\kappa_s k_{off} v_u}{n_b\kappa_c}, \qquad t < t_{rupt} \tag{S10}$$

We address the motor-dominated regime and the intermediate regime separately.

*Motor-dominated regime*

Motor-dominated protrusions operate at optimum conditions when most of the elastic energy in the clutches-substrate axis is taken up at the beginning of the cell cycle by the substrate. Mathematically, this can be expressed as $\kappa_s \gg \kappa_c$. Under these conditions, the substrate deforms at a rate approximately equal to the retrograde flow, the unloaded myosin velocity: $dx_s/dt \sim v_u$ (see Eq. (S17)). The instantaneous traction forces produced at time $t$ therefore scale as $T \sim v_u\kappa_s t$. This implies that first-bound clutch extension evolves as $x_c \sim (v_u\kappa_s/\kappa_c)t$ (see Eq. (S82)), and reaches an extension value equal to its characteristic rupture length $\ell_{rupt} = F_b/\kappa_c$ at a time $t_{rupt}^{single} \sim F_b/v_u\kappa_s$. Notice that the clutch rupture time is independent of clutch stiffness since both rupture length and clutch elongation rate scale inversely to clutch stiffness. The effective clutch binding rate is initially $t_{bind}^{single} \sim 1/n_c k_{on}$. We can determine an upper bound limit of the optimum substrate stiffness that maximizes traction forces $\kappa_s^{opt}$ by realizing that the necessary condition for the protrusion to operate at its optimum is that clutch binding rates are faster than clutch rupture rates: $t_{rupt}^{single} > t_{bind}^{single}$. Thus, $\kappa_s^{opt} < n_c F_b k_{on}/v_u$. We now proceed to estimate $\kappa_s^{opt}$. A careful scaling analysis of Eq. (S10) suggests that in the limit of high motor activity, the clutch extension rate scales as $v_{e_c} \sim \kappa_s v_u (k_{on} + k_{off})/n_c \kappa_c k_{on}$ where we have used Eq. (S6). An increase in the number of available clutches $n_c$, the clutch stiffness constant $\kappa_c$, or the ratio $k_{on}/k_{off}$ reinforces clutches augmenting the clutch resistance against myosin pulling forces, resulting in lower clutch deformation rates. Also, an increase in the load-free velocity of myosin motors results in stronger actin retrograde flows, giving rise to higher clutch extension rates. Because softer substrates undergo higher deformations than more rigid substrates, lower clutch extension rates are expected for more compliant substrates, as predicted by scaling analysis and in agreement with previous studies (3,4). Our scaling for $v_{e_c}$ implies that the clutch rupture time $t_{rupt}$ scales as $t_{rupt} \sim k_{on} n_c F_b/\kappa_s v_u (k_{on} + k_{off})$. We apply the optimal condition $t_{bind} \sim t_{rupt}$ to determine the substrate stiffness that maximizes traction forces in a motor-dominated protrusion

$$\kappa_{s,m}^{opt} = C_2 \frac{n_c F_b k_{on}}{v_u}, \qquad \text{(S11)}$$

Our scaling analysis allows us to obtain the optimum substrate stiffness up to an unknown constant $C_1$. The upper limit calculated earlier for $\kappa_{s,m}^{opt}$ indicates that $C_2 < 1$. We estimate the constant $C_2$ by fitting Eq. (27) to our numerical results. We get $C_2 = 0.4$, consistent with our upper limit calculation. We find very good agreement between our theoretical prediction (Eq. (S11)), the numerical solution of the mean-field model and the solution of the dimensionless version of the stochastic motor-clutch model (3), as shown in Fig. (S3, right). According to Eq. (S11), the optimum substrate stiffness is proportional to the characteristic maximum clutch elastic force $n_c F_b$ and inversely proportional to the distance that unloaded actin filaments translocate within the characteristic clutch binding time $v_u/k_{on}$. Our theoretical solution also suggests that in the high-motor regime, the optimum substrate stiffness is myosin-insensitive and independent of clutch stiffness $\kappa_c$.

*Motor-clutch balanced regime*

We now obtain an expression for the optimum substrate stiffness that maximizes traction forces for protrusions that lie in the intermediate balanced regime. As clutches get stronger either by an increase in the clutch elastic force capacity $n_c F_b$ or an increase in the ratio $k_{on}/k_{off}$, and/or myosin motors get weaker by a reduction in the total myosin stall force $n_m F_m$, time-averaged actin retrograde flows get weaker, and $t_{rupt}$ increases. This rise in clutch rupture time can be compensated by an increase in substrate stiffness, so that optimum conditions for maximum traction force $t_{bind} \sim t_{rupt}$ are met. Therefore, we expect that as clutches become dominant over motors, the optimum substrate stiffness for traction force will shift towards higher stiffnesses, as previously reported (4,5). We indeed observe this in Fig (4) as well as in Eq. (6). We go ahead and estimate $t_{rupt}$ from Eq. (S10). Before clutches rupture under load, the clutch strain rate scales as $v_e \sim \kappa_s k_{off} v_u \Big/ \left[k_{on}(n_c - n_b)\kappa_c + \frac{n_b \kappa_c v_u \kappa_s}{n_m F_m}\right]$. The characteristic clutch rupture time $t_{rupt} = F_b/\kappa_c v_{e_c}$ scales as $t_{rupt} \sim \frac{k_{on}}{k_{on}+k_{off}} n_c F_b \frac{1}{\kappa_s v_u}\left(1 + C_3 \frac{\kappa_s v_u}{k_{off} n_m F_m}\right)$, where $C_3$ is an unknown parameter. Notice that our scaling analysis does not allow us to determine this unknown parameter analytically. We apply the condition $t_{bind} \sim t_{rupt}$ to determine the substrate stiffness that maximizes traction forces in a motor-clutch balanced protrusion:

$$\kappa_{s,i}^{opt} = C_2 \frac{n_c F_b k_{on}}{v_u} \frac{1}{1 - C_3 \dfrac{k_{on}}{k_{off}} \dfrac{n_c F_b}{n_m F_m}}, \qquad (S12)$$

which corresponds with Eq. (6) in the main text. The value of $C_3$ is obtained by fitting Eq. (S12) with our numerical results. We get $C_3 = 0.05$. Figures (1F) and (S3) show that our theoretical expression is in very good agreement with our numerical results for all the motor-clutch dimensionless parameters. The optimum substrate stiffness in a protrusion that belongs to the intermediate regime, $\kappa_{s,i}^{opt}$, is myosin sensitive and independent of the effective clutch stiffness $\kappa_c$. The optimum substrate stiffness in the motor-dominated regime $\kappa_{s,m}^{opt}$ (Eq. (S11)) is recovered in Eq. (S12) by taking the limit $C_3 k_{on} n_c F_b / k_{off} n_m F_m \to 0$. The optimum stiffness for maximum traction force has been previously estimated and left as a function of an unknown parameter $\varepsilon$, the fraction of the theoretical maximum load that a protrusion can generate (6). This unknown parameter depends on some of the motor-clutch parameters. Equating Eq. (30) in reference (6) with Eq. (S12) in the current study, we find $\varepsilon = e^{-\ln n_c/(F - C_3\tau)}$.

**Critical motor activity parameter that sets the boundary between protrusions in the stalled regime and balanced regime on soft substrates**

The boundary between the clutch-dominated stalled regime and the balanced regime is set by the condition $t_s^{max} \sim t_{rupt}$ — that is, the time required to reach stall conditions matches the time needed for clutch bonds to break due to force. The time-evolution of the substrate deformation rate obeys the following ordinary differential equation:

$$\frac{dx_s}{dt} = \frac{\kappa_c}{\kappa_s + n_b\kappa_c}\left[v_u n_b\left(1 - \frac{\kappa_c \ell_b}{n_m F_m}\right) - k_{off}\int_{-\infty}^{\infty} x_c e^{\kappa_c |x_c|/F_b} P_b dx_c\right]. \quad (S13)$$

To estimate the time required for the substrate to reach its maximum deformation $t_s^{max}$, we approximate the time-evolution equation of the substrate deformation rate for times shorter than the clutch rupture time

$$\frac{dx_s}{dt} \approx \frac{n_b\kappa_c}{\kappa_s + n_b\kappa_c} v_u\left[1 - \left(1 + \frac{n_m F_m k_{off}}{n_b \kappa_c v_u}\right)\frac{x_s}{x_s^{max}}\right] \quad t < t_{rupt} \qquad (S14)$$

where the maximum substrate deformation is $x_s^{max} = n_m F_m/\kappa_s$. After a small transient from the onset of loading, we expect that in the clutch dominated regime $n_b \kappa_c v_u \gg n_m F_m k_{off}$. Under this assumption, Eq. (S14) reduces to

$$\frac{dx_s}{dt} \approx \frac{n_b \kappa_c}{\kappa_s + n_b \kappa_c} v_u \left(1 - \frac{x_s}{x_s^{max}}\right), \quad t < t_{rupt} \tag{S15}$$

which suggests that the time to reach maximum substrate deformations $t_s^{max}$ scales as

$$t_s^{max} \sim \frac{\kappa_s + n_b \kappa_c}{n_b \kappa_c} \frac{n_m F_m}{\kappa_s v_u}. \tag{S16}$$

We expect that, after a small transient from the onset of loading, the clutch ensemble stiffness is much more rigid than the substrate stiffness, i.e. $n_b \kappa_c \gg \kappa_s$. We expect this condition to be satisfied if the substrate is soft enough, and sufficient number of clutches are bound. Under this assumption, Eqs. (S15) and (S16) simplify to

$$\frac{dx_s}{dt} \approx v_u \left(1 - \frac{x_s}{x_s^{max}}\right), \quad t < t_{rupt} \tag{S17}$$

$$t_s^{max} \sim \frac{n_m F_m}{\kappa_s v_u}. \tag{S18}$$

We can think of clutches and substrate as two mechanically connected entities, with clutch ensemble stiffness and substrate stiffness $n_b \kappa_c$ and $\kappa_s$, respectively. At the time when $n_b \kappa_c \gg \kappa_s$, substrate deformation is much greater than the deformation that any individual clutch undergoes. Consequently, substrates deform at a rate equal to the F-actin retrograde velocity $v_{act}$, as indicated by Eq. (S17). As long as the system is far away from stall conditions, i.e. $x_s \ll x_s^{max}$, the substrate strain rate is approximately equal to the myosin load-free velocity. The timescale to reach maximum substrate deformations then scales as $t_s^{max} \sim x_s^{max}/v_u$. The transition between the clutch-dominated regime and the intermediate regime will occur when $t_s^{max} \sim t_{rupt}$ — that is,

$$\frac{n_m F_m}{n_c F_b} \sim \frac{k_{on}}{k_{on} + k_{off}}, \tag{S19}$$

where we have assumed that $n_b \kappa_c \gg \kappa_s$ after a short period of time after the beginning of loading. Therefore, the protrusion will be in the clutch-dominated regime when

$$F < F_{c-i}^{crit} = C_1 \frac{k_{on}}{k_{on} + k_{off}}, \tag{S20}$$

where $C_1$ is a constant that we estimate by fitting Eq. (S20) with our numerical results. We find $C_1 = 0.4$.

**Derivation of the mean traction force, number of bound clutches and clutch elongation rates of protrusions on rigid substrates**

In our work, we have used two approaches to study the dynamics of cell protrusions: a stochastic Langevin-type approach (3) and a mean-field approach. Whereas the stochastic approach is suitable to a very small timescale, on which stochastic fluctuations in traction forces are observed, the mean-field model addresses a much coarser timescale. On rigid substrates, cycles of loading/unloading occur at a very high frequency. Because the frequency of load-and-fail dynamics is so high, the load-and-fail cycling time is smaller than the minimum timescale addressed by the mean-field model. As a result, the mean-field framework is not able to capture these periodic events on rigid substrates, and its temporal solution reaches steady-state after the first loading event. This allows us to determine the time-averaged traction force $\overline{T}$, mean number of bound clutches $\bar{n}_b$ and mean clutch strain rate $\bar{v}_e$ in a more theoretical way by seeking the steady-state solution of the density conservation equation:

$$\frac{d\overline{P}_b}{dx_c} = \frac{k_{on}(n_c - \bar{n}_b)}{\bar{v}_e}\delta(x_c) - \frac{k_{off}}{\bar{v}_e}\, e^{\frac{\kappa_c|x_c|}{F_b}}\overline{P}_b \qquad \text{(S21)}$$

where the mean clutch strain rate $\bar{v}_e$ reads

$$\bar{v}_e = v_u\left(1 - \frac{\kappa_c}{n_m F_m}\int_{-\infty}^{\infty} x_c P_b(x_c, t)dx_c\right) \qquad \text{(S22)}$$

Integration of Eq. (S21) over a small region $x_c\,\epsilon[-\varepsilon, \varepsilon]$ and taking the limit $\varepsilon \to 0$ yields

$$\overline{P}_b(0^+) = \frac{k_{on}(n_c - \bar{n}_b)}{\bar{v}_e} \qquad \text{(S23)}$$

where we have assumed that $\overline{P}_b(0^-) \to 0$. We can easily seek a solution for the mean probability density for $x_c > 0$ by applying separation of variables to Eq. (S21) and making use of Eq. (S23) to solve for the integration constant. We find that $\overline{P}_b$ has a double exponential functional form with a very fast decay at a clutch length equal to the clutch rupture length $\ell_{rupt} = F_b/\kappa_c$:

$$\overline{P}_b(x_c) = \frac{k_{on}(n_c - \overline{n}_b)}{\overline{v}_e} e^{\frac{F_b k_{off}}{\kappa_c \overline{v}_e}\left(1 - e^{\frac{\kappa_c}{F_b} x_c}\right)} \tag{S24}$$

Here, $\overline{v}_e$ is still unknown. Taking the zeroth-order moment of Eq. (S24) and rearranging terms, we find

$$\overline{n}_b = \frac{k_{on}\beta}{k_{on}\beta + \overline{v}_e} n_c \tag{S25}$$

where

$$\beta = \frac{F_b}{\kappa_c} e^{\frac{F_b k_{off}}{\kappa_c \overline{v}_e}} \Gamma\left(0, \frac{F_b k_{off}}{\kappa_c \overline{v}_e}\right) \tag{S26}$$

Here, $\Gamma$ is the incomplete gamma function. Inserting Eq. (S25) into Eq. (S24) yields

$$\overline{P}_b(x_c) = \frac{k_{on}}{k_{on}\beta + \overline{v}_e} n_c e^{\frac{F_b k_{off}}{\kappa_c \overline{v}_e}\left(1 - e^{\frac{\kappa_c}{F_b} x_c}\right)} \tag{S27}$$

We plug Eq. (S27) into Eq. (S21) to obtain a non-linear equation for the mean clutch extension rate

$$\overline{v}_e{}^2 + (k_{on}\beta - v_u)\overline{v}_e + v_u k_{on}\left(\frac{n_c \kappa_c \alpha}{n_m F_m} - \beta\right) = 0 \tag{S28}$$

where

$$\alpha = \left(\frac{F_b}{\kappa_c}\right)^2 e^{\frac{F_b k_{off}}{\kappa_c \overline{v}_e}} G_{2\,3}^{3\,0}\left(\frac{F_b k_{off}}{\kappa_c \overline{v}_e}\middle| \begin{matrix} 1 & 1 & \\ 0 & 0 & 0 \end{matrix}\right) \tag{S29}$$

Here, $G$ is the Meijer G-function. We can numerically solve Eq. (S28) to determine $\overline{v}_e$. Once we determine $\overline{v}_e$, we can compute $\overline{n}_b$ and $\overline{P}_b(x_c)$ using Eqs. (S25) and (S27), respectively. The mean traction force can be then obtained as

$$\overline{T} = \frac{G_{2\,3}^{3\,0}\left(\frac{F_b k_{off}}{\kappa_c \overline{v}_e}\middle| \begin{matrix} 1 & 1 & \\ 0 & 0 & 0 \end{matrix}\right)}{\Gamma\left(0, \frac{F_b k_{off}}{\kappa_c \overline{v}_e}\right)} \overline{n}_b F_b, \tag{S30}$$

Equation (S30) shows that the mean traction force produced by an individual protrusion on a rigid substrate is proportional to the mean clutch elastic force $\bar{n}_b F_b$ multiplied by a pre-factor that depends on $F_b k_{off}/\kappa_c \bar{v}_e$. We rewrite Eqs. (S24−S30) in dimensionless form:

$$\bar{n}_b^* = \frac{\tau\beta^*}{\tau\beta^* + \bar{v}_e^*}, \qquad \beta^* = \frac{1}{F}\int_0^\infty e^{\frac{1}{F\bar{v}_e^*}(1-e^t)} dt, \qquad \bar{P}_b^*(x_c^*) = \frac{\bar{n}_b^*}{\beta^*} e^{\frac{1}{F\bar{v}_e^*}\left(1-e^{Fx_c^*}\right)} \qquad \text{(S31 − S33)}$$

$$\bar{v}_e^{*2} + \left(\tau\beta^* - \frac{\omega}{F}\right)\bar{v}_e^* + \frac{\tau\omega}{F}(\alpha^* - \beta^*) = 0, \qquad \alpha^* = \frac{1}{F^2}\int_0^\infty t e^{\frac{1}{F\bar{v}_e^*}(1-e^t)} dt, \qquad \bar{T}^* = F\,\bar{n}_b^* \frac{\alpha^*}{\beta^*} \quad \text{(S34 − S36)}$$

We solve for $\bar{v}_e^*$ by numerically solving the nonlinear equation (S34). Once we solve for the mean clutch extension rate, we can compute $\bar{n}_b^*$ and $\bar{T}^*$ using Eqs. (S31) and (S36), respectively. We now proceed to determine analytical expressions for two asymptotic cases: low-motor activity ($F \to 0$) and high-motor activity ($F \to \infty$).

*Low-motor activity ($F \to 0$)*

We first derive an analytical expression for the dimensionless clutch strain rate $\bar{v}_e^*$. We use regular perturbation theory and expand $\bar{v}_e^*$ in powers of F:

$$\bar{v}_e^* = v_0 + F v_1 + F^2 v_2 + \mathcal{O}(F^3) \qquad \text{(S37)}$$

We substitute the assumed perturbation series into Eqs. (S32) and (S33), and after some math we get:

$$\beta^* = v_0 + F(v_1 - v_0^2) + F^2(2v_0^3 - 2v_0 v_1 + v_2) + \mathcal{O}(F^3) \qquad \text{(S38)}$$

$$\alpha^* = v_0^2 + F(2v_0 v_1 - 3v_0^3) + F^2(v_1^2 + 2v_0 v_2 - 9v_0^2 v_1 + 11v_0^4) + \mathcal{O}(F^3) \qquad \text{(S39)}$$

We insert Eqs. (S37-S39) into Eq. (S34), and find that the first three leading order terms in the expansion satisfy the following equations:

$$F_0: \qquad \tau(v_0 - 1) - 1 = 0, \qquad \text{(S40)}$$

$$F_1: \qquad (\tau + 1)v_0^2 - \omega v_1 + \tau\omega(-3v_0^3 + v_0^2 + 2v_0 v_1 - v_1) = 0, \qquad \text{(S41)}$$

$$\mathrm{F}_2: \quad (2+\tau)v_0v_1 + \tau v_0(v_1 - v_0^2) - \omega v_2 + \tau\omega(v_1^2 + 2v_0v_2 - 9v_0^2v_1 + 11v_0^4) +$$

$$-\tau\omega(2v_0^3 - 2v_0v_1 + v_2) = 0. \tag{S42}$$

The leading-order solution can be easily obtained from Eq. (S40):

$$v_0 = \frac{\tau+1}{\tau} \tag{S43}$$

We recursively solve for higher order terms by solving Eqs. (S41) and (S42), we get:

$$v_1 = \left(\frac{\tau+1}{\tau}\right)^2\left(3 - \frac{1}{\omega}\right) - \frac{\tau+1}{\tau} \tag{S44}$$

$$v_2 = \frac{\tau+1}{\tau^3\omega^2}[1 - 9\omega + 7\omega^2 + \tau^2(1 - 6\omega + \omega^2) + \tau(2 - 15\omega + 7\omega^2)] \tag{S45}$$

An expression for the dimensionless time-averaged clutch elongation rate can then be obtained by substituting Eqs. (S43-S45) into the perturbation expansion in Eq. (S37):

$$\bar{v}_e^* = \frac{\tau+1}{\tau} + \left(\frac{\tau+1}{\tau}\right)^2\left(\frac{2\tau+3}{\tau+1} - \frac{1}{\omega}\right)\mathrm{F} +$$

$$+\frac{\tau+1}{\tau^3\omega^2}[1 - 9\omega + 7\omega^2 + \tau^2(1 - 6\omega + \omega^2) + \tau(2 - 15\omega + 7\omega^2)]\mathrm{F}^2 + \mathcal{O}(\mathrm{F}^3) \tag{S46}$$

Eq. (S43) agrees very well with our numerical solutions as shown in the insets in Fig. (3). In dimensional form, the time-averaged clutch elongation rate reads

$$\bar{v}_e \approx \frac{n_m F_m k_{off}}{n_c \kappa_c}\left(1 + \frac{k_{off}}{k_{on}}\right)\left[1 + \frac{n_m F_m}{n_c F_b}\left(\left(1 + \frac{k_{off}}{k_{on}}\right)\left(3 - \frac{F_b k_{off}}{v_u \kappa_c}\right) - 1\right)\right] \tag{S47}$$

where we have only kept the first two leading order terms, for simplicity. Equation (S47) indicates that an increase in myosin forces strengthens rearward actin flows, in turn enhancing mean clutch elongation rates. It also indicates that an increase in the total number of clutches $n_c$, the clutch stiffness $\kappa_c$, the clutch association constant $k_{on}$, or the characteristic clutch rupture force $F_b$ decreases averaged clutch elongation rates by strengthening clutches, whereas an increase in the myosin load-free velocity $v_u$ or in the unloaded clutch dissociation constant $k_{off}$ increases clutch extension rates. Because only large values of $\omega$ are physiologically relevant, we can simplify Eq. (S47) by taking the limit for large $\omega$. We get,

$$\bar{v}_e^*(\omega \to \infty) = \frac{\tau+1}{\tau}\left[1 + \frac{2\tau+3}{\tau}F + \frac{7+7\tau+\tau^2}{\tau^2}F^2\right] + \mathcal{O}(F^3) \tag{S48}$$

Next, we seek a solution for the time-averaged fraction of bound clutches. We substitute the results obtained in Eqs. (S43−S45) into Eqs. (S38) and (S39), and plug the results for $\beta^*$ and $\alpha^*$ along with Eq. (S46) into Eq. (S31); upon simplification, we get:

$$\bar{n}_b^* = \frac{\tau}{\tau+1} - \frac{1}{\tau+1}F + \frac{1-\omega}{\tau\omega}F^2 + \mathcal{O}(F^3) \tag{S49}$$

Very good agreement is found between Eq. (S49) and our numerical results, as shown in Fig. (3, middle). In dimensional form, the time-averaged number of bound clutches reads

$$\bar{n}_b \approx \frac{k_{on}}{k_{on}+k_{off}}\left(n_c - \frac{n_m F_m}{F_b}\frac{k_{off}}{k_{on}} + \frac{(n_m F_m)^2}{n_c F_b^2}\right) \tag{S50}$$

where we have only kept the first two leading order terms, for simplicity. The leading order term in Eq. (S50) corresponds to the equilibrium number of bound clutches in an unloaded state, set by just a balance of binding/unbinding kinetics. As expected, an increase in myosin forces reduces the time-averaged number of bound clutches, as indicated by the negative sign in front of the second term in Eq. (S50). The first two leading order terms do not depend on the parameter $\omega$. Therefore, we must look at the $F^2$ term in Eq. (S49) to study the dependence of number of bound clutches on clutch stiffness $\kappa_c$ and myosin-load free velocity $v_u$. We find that larger values of $\kappa_c$ or $v_u$ (larger $\omega$) leads to a reduction in the average number of bound clutches, consistent with our discussion of Fig. (2) in the main text and with Fig. (S4, right). Finally, we seek a formula for the time-averaged traction force. We substitute Eqs. (S38), (S39) and (S49) into Eq. (S36) and simplify, to get

$$\bar{T}^* = F - \frac{\tau+1}{\tau\omega}F^2 + \frac{1+2\tau+\tau^2-\omega(3+5\tau+2\tau^2)}{\omega^2\tau^2}F^3 + \mathcal{O}(F^4) \tag{S51}$$

In dimensional form, the time-averaged traction force reads

$$\overline{T} \approx n_m F_m \left[1 - \frac{n_m F_m k_{off}^2}{n_c \kappa_c v_u k_{on}} \left(1 + \frac{k_{on}}{k_{off}} + \frac{\kappa_c v_u}{k_{on} F_b} \left(1 + 2\frac{k_{on}}{k_{off}}\right)\right)\right] \tag{S52}$$

where we have only kept the first two leading order terms, for simplicity.

*High-motor activity ($F \to \infty$)*

We proceed now to obtain analytical expressions for $\overline{T}$, $\bar{n}_b$ and $\bar{v}_e$ for motor-dominated protrusions. At high motor activity, time-averaged traction force and number of bound clutches are independent of motor activity, as demonstrated by the plateau in Figs. (3A) and (3B) for large $F$. We take the limit $F \to \infty$ in Eq. (S34) and look for the leading-order term, we get

$$(\bar{v}_e^* + \tau\beta^*)\left(\bar{v}_e^* - \frac{\omega}{F}\right) \to 0. \tag{S53}$$

Only positive values of $\bar{v}_e^*$ are physically possible, thus solution of Eq. (S53) reads

$$\bar{v}_e^* \to \frac{\omega}{F} \tag{S54}$$

Therefore, clutches of motor-dominated protrusions elongate on rigid substrates at an average rate that approaches the myosin load-free velocity $\bar{v}_e = v_u$. The mean number of bound clutches of a motor-dominated protrusion can then be obtained by plugging Eq. (S32) into Eq. (S31) and making use of Eq. (S54), we get:

$$\bar{n}_b^* \to \frac{\tau e^{\frac{1}{\omega}} \Gamma\left(0, \frac{1}{\omega}\right)}{\tau e^{\frac{1}{\omega}} \Gamma\left(0, \frac{1}{\omega}\right) + \omega} \tag{S55}$$

Equation (S55) is in very good agreement with the stochastic model results, as shown in Fig. (4C). The fraction of bound clutches monotonically increases as the clutch kinetic parameter $\tau$ increases and/or the parameter $\omega$ decreases. In dimensional form, Eq. (S55) reads

$$\bar{n}_b \to \frac{k_{on} F_b e^{\frac{F_b k_{off}}{\kappa_c v_u}} \Gamma\left(0, \frac{F_b k_{off}}{\kappa_c v_u}\right)}{k_{on} F_b e^{\frac{F_b k_{off}}{\kappa_c v_u}} \Gamma\left(0, \frac{F_b k_{off}}{\kappa_c v_u}\right) + v_u \kappa_c} n_c, \tag{S56}$$

Finally, we can easily obtain the time-averaged force transmitted to the substrate:

$$\bar{T}^* \to \frac{G_{2\,3}^{3\,0}\left(\frac{1}{\omega}\middle|\begin{matrix}1 & 1 & \\ 0 & 0 & 0\end{matrix}\right)}{\Gamma\left(0,\frac{1}{\omega}\right)}\bar{n}_b^* \tag{S57}$$

In dimensional form, Eq. (S57) reads

$$\bar{T} \to \frac{G_{2\,3}^{3\,0}\left(\frac{F_b k_{off}}{\kappa_c v_u}\middle|\begin{matrix}1 & 1 & \\ 0 & 0 & 0\end{matrix}\right)}{\Gamma\left(0,\frac{F_b k_{off}}{\kappa_c v_u}\right)}\bar{n}_b F_b, \tag{S58}$$

where $\bar{n}_b$ is that in Eq. (S56).

**Derivation of the mean traction force produced by cell protrusions with reinforcement on rigid substrates**

The frequency of load-and-fail dynamics is so high on rigid substrates that the mean-field framework cannot capture these periodic events and the mean-field temporal solution reaches steady-state after the first loading event. This steady-solution corresponds to the time-averaged solution of the stochastic motor-clutch model, allowing us to determine the time-averaged traction force $\bar{T}$, mean number of bound clutches $\bar{n}_b$ and mean clutch strain rate $\bar{v}_e$ in a more theoretical way by seeking the steady-state solution of Eq. (76). The time-averaged probability density of clutch extensions for a protrusion with clutch reinforcement has also a double exponential form on clutch extensions:

$$\bar{P}_b(x_c) = \frac{k_{on}\left(1+\theta\frac{\bar{n}_b}{n_c}\right)(n_c-\bar{n}_b)}{\bar{v}_e} e^{\frac{F_b k_{off}}{\kappa_c \bar{v}_e}\left(1-e^{\frac{\kappa_c}{F_b}x_c}\right)}. \tag{S59}$$

where we have assumed in the subsequent derivation that the clutch reinforcement threshold force is $F_{th} = 0$. The mean clutch strain rate and mean number of bound clutches take the form:

$$\bar{v}_e = v_u\left[1-\frac{\kappa_c k_{on}}{n_m F_m \bar{v}_e}\left(1+\theta\frac{\bar{n}_b}{n_c}\right)(n_c-\bar{n}_b)\sigma\right] \tag{S60}$$

and

$$\bar{n}_b = \frac{k_{on}(1 + \theta\frac{\bar{n}_b}{n_c})(n_c - \bar{n}_b)}{\bar{v}_e}\rho \qquad (S61)$$

where

$$\sigma = \int_0^\infty e^{\frac{F_b k_{off}}{\kappa_c \bar{v}_e}\left(1 - e^{\frac{\kappa_c}{F_b}x_c}\right)} x_c dx_c, \qquad (S62)$$

and

$$\rho = \int_0^\infty e^{\frac{F_b k_{off}}{\kappa_c \bar{v}_e}\left(1 - e^{\frac{\kappa_c}{F_b}x_c}\right)} dx_c \qquad (S63)$$

We can express $\bar{n}_b$ as function of $\bar{v}_e$ by rearranging terms in Eq. (79):

$$\bar{n}_b = \frac{n_c}{2\theta}\left[-\left(\frac{\bar{v}_e}{k_{on}\rho} + 1 - \theta\right) + \sqrt{\left(\frac{\bar{v}_e}{k_{on}\rho} + 1 - \theta\right)^2 + 4\theta}\right] \qquad (S64)$$

We plug Eq. (82) into Eq. (78) and obtain a nonlinear integral equation for the mean clutch extension rate:

$$1 - \frac{\bar{v}_e}{v_u} + \frac{\kappa_c \sigma n_c}{2\theta n_m F_m \rho}\left[\left(\frac{\bar{v}_e}{k_{on}\rho} + 1 - \theta\right) - \sqrt{\left(\frac{\bar{v}_e}{k_{on}\rho} + 1 - \theta\right)^2 + 4\theta}\right] = 0 \qquad (S65)$$

The traction force produced by the protrusion can then be expressed as

$$\bar{T} = \frac{\kappa_c \bar{n}_b \sigma}{\rho} \qquad (S66)$$

In dimensionless form, Eqs. (82-84) read:

$$\bar{n}_b^* = \frac{1}{2\theta}\left[-\left(\frac{\bar{v}_e^*}{\tau\rho^*} + 1 - \theta\right) + \sqrt{\left(\frac{\bar{v}_e^*}{\tau\rho^*} + 1 - \theta\right)^2 + 4\theta}\right] \qquad (S67)$$

$$4\theta^2 \rho^{*2}\left(1 - \frac{F}{\omega}\bar{v}_e^*\right)^2 + 4\theta\sigma^*\left[\left(1 - \frac{F}{\omega}\bar{v}_e^*\right)\left(\frac{\bar{v}_e^*}{\tau} + \rho^*(1 - \theta)\right) - \sigma^*\right] = 0 \qquad (S68)$$

$$\bar{T}^* = \frac{F\bar{n}_b^* \sigma^*}{\rho^*} \qquad (S69)$$

where $\rho^* = \rho n_c \kappa_c / n_m F_m$ and $\sigma^* = \sigma n_c^2 \kappa_c^2 / n_m^2 F_m^2$.

We further proceed to obtain asymptotic expressions for the time-averaged traction force and mean number of bound clutches for motor-dominated protrusions with reinforcement. The leading order term can be obtained by taking the limit $\mathrm{F} \to \infty$ on Eqs. (77−84). After some math, we get:

$$\bar{n}_b^* \to \frac{-\left(\frac{\omega}{\tau e^{1/\omega}\Gamma(0,1/\omega)} + 1 - \theta\right) + \sqrt{\left(\frac{\omega}{\tau e^{1/\omega}\Gamma(0,1/\omega)} + 1 - \theta\right)^2 + 4\theta}}{2\theta} \tag{S70}$$

$$\bar{T}^* \to \frac{G_{2\,3}^{3\,0}\left(\frac{1}{\omega}\middle|\begin{matrix}1 & 1 & \\ 0 & 0 & 0\end{matrix}\right)}{\Gamma\left(0,\frac{1}{\omega}\right)}\bar{n}_b^* \tag{S71}$$

$$\bar{P}_b^*(x_c^*) \to \frac{F\tau}{\omega}\left(1 + \theta\frac{\bar{n}_b^*}{n_c}\right)(1 - \bar{n}_b^*)e^{\frac{1-e^{Fx_c^*}}{\omega}} \tag{S72}$$

We additionally take the limit of Eq. (S70) when $\theta \to 0$ to explore traction force generation on protrusions with low reinforcement. We get:

$$\bar{n}_b^* \to \frac{1}{1 + \frac{\omega}{\tau e^{1/\omega}\Gamma(0,1/\omega)}} + \frac{\frac{\omega}{\tau e^{1/\omega}\Gamma(0,1/\omega)}}{\left(1 + \frac{\omega}{\tau e^{1/\omega}\Gamma(0,1/\omega)}\right)^3}\theta \tag{S73}$$

where we have only kept the first two leading order terms in $\theta$, for simplicity. The positive sign in front of the last term in Eq. (S73) indicates that reinforcement enhances the mean number of bound clutches and, therefore, the mean traction force produced by the protrusion as shown in Figs. (6E) and (6F).

**Dynamics of strong-motor protrusions are myosin independent**

In this section, we demonstrate using a simple approach that clutch dynamics of strong-motor protrusions do not depend on myosin activity. We explore the evolution of an individual protrusion right after the first clutch connects the actin-cytoskeleton with the surrounding substrate. Before any clutch couples the actin network with the substrate, the actin cytoskeleton flows rearwards at the myosin load-free velocity. Once the first clutch binds, force balance reads

$$n_m F_m \left(1 - \frac{v_{act}}{v_u}\right) = \kappa_c x_c = \kappa_s x_s. \tag{S74}$$

The clutch extension rate is equal to the difference between actin retrograde flow velocity and the substrate deformation rate:

$$\frac{dx_c}{dt} = v_{act} - \frac{dx_s}{dt}. \tag{S75}$$

Combination of Eqs. (S6) and (S7) yields an ordinary differential equation for the actin retrograde flow velocity:

$$\frac{n_m F_m}{v_u}\left(\frac{1}{\kappa_c} + \frac{1}{\kappa_s}\right)\frac{dv_{act}}{dt} = -v_{act}. \tag{S76}$$

We easily solve for Eq. (S8) by applying separation of variables:

$$v_{act}(t) = v_u e^{-\frac{v_u}{n_m F_m\left(\frac{1}{\kappa_c}+\frac{1}{\kappa_s}\right)}t} \tag{S77}$$

We substitute Eq. (S9) into Eq. (S6) to obtain:

$$x_c(t) = \frac{\kappa_s}{\kappa_c} x_s(t) = \frac{n_m F_m}{\kappa_c}\left(1 - e^{-\frac{v_u}{n_m F_m\left(\frac{1}{\kappa_c}+\frac{1}{\kappa_s}\right)}t}\right) \tag{S78}$$

If the bound clutch does not dissociate before reaching its characteristic rupture length $x_c^{rupt} = F_b/\kappa_c$ and that no other clutch binds, Eq. (S10) implies that the time required for the clutch to reach its rupture length $t_r$ is:

$$t_r = -\frac{n_m F_m}{v_u}\left(\frac{1}{\kappa_c} + \frac{1}{\kappa_s}\right)\ln\left(1 - \frac{F_b}{n_m F_m}\right). \tag{S79}$$

Notice that the argument of the logarithm in Eq. (S11) is positive provided that the characteristic clutch rupture length $x_c^{rupt}$ is smaller than the clutch elongation at stall conditions $x_c^{stall} = n_m F_m/\kappa_c$. Therefore, Eq. (S11) is valid for $x_c^{rupt} < x_c^{stall}$. We take the high-myosin limit ($n_m F_m \to \infty$) of Eqs. (S10) and (S11) to obtain

$$\lim_{n_m F_m \to \infty} x_c(t) = \frac{v_u t}{1 + \kappa_c/\kappa_s}\left[1 - \frac{v_u t}{2 n_m F_m \left(\frac{1}{\kappa_c} + \frac{1}{\kappa_s}\right)}\right] + \mathcal{O}\left(\frac{t^3}{(n_m F_m)^2}\right), \tag{S80}$$

$$\lim_{n_m F_m \to \infty} t_r = \frac{F_b}{v_u}\left(\frac{1}{\kappa_c} + \frac{1}{\kappa_s}\right) + \mathcal{O}\left(\frac{1}{n_m F_m}\right). \tag{S81}$$

One can clear see that, in the high-myosin limit and for times on the order of the clutch rupture time $t = \mathcal{O}(t_r)$, clutch elongation is asymptotically myosin independent. Consequently, the time required for the clutch to reach its rupture length is also myosin independent, as Eq. (S13) indicates. Hence, the clutch elongation rate is time independent and asymptotically equal to

$$\lim_{n_m F_m \to \infty} \frac{dx_c(t)}{dt} = \frac{v_u}{1 + \kappa_c/\kappa_s}. \tag{S82}$$

We use Eqs. (S6) and (S14) to obtain the asymptotic substrate deformation rate:

$$\lim_{n_m F_m \to \infty} \frac{dx_s(t)}{dt} = \frac{v_u}{1 + \kappa_s/\kappa_c}, \tag{S83}$$

which indicates that the dynamics of motor-dominated protrusions are myosin insensitive, in agreement with our numerical and analytical results. Notice that the average time required for a second clutch to bind is equal to $1/(n_c - 1)k_{on}$. A second clutch will on average bind before the first bound clutch reaches its rupture length when

$$\frac{F_b k_{on}(n_c - 1)}{v_u}\left(\frac{1}{\kappa_c} + \frac{1}{\kappa_s}\right) > 1. \tag{S84}$$

**Numerical solution of the mean-field conservation equation**

We solve Eqs. (14-16) of the main text numerically for a range of parameter values. Note that the probability density $P_b^*(x_c^*, t^*)$ is defined on an infinite domain on clutch extensions. We use a sufficiently large computational domain so that domain size does not affect the numerical solution. We impose that the probability density vanishes at the two boundaries, i.e. $P_b^*(\pm\infty, t^*) = 0$. We discretize the domain using a uniform grid, approximate integrals using the trapezoidal quadrature method, and approximate the first derivative in Eq. (14) using the first order left-sided finite difference scheme to guarantee numerical stability. We approximate the Dirac delta function with the following function:

$$\delta^*(x_c^*) \approx \frac{1}{\pi}\frac{\varepsilon}{x_c^{*2} + \varepsilon^2}, \tag{S85}$$

where we have used $\varepsilon = 5 \times 10^{-3}$ in all simulations. We have integrated Eq. (14) in time by using the Forward-Backward Euler method, where we have treated the clutch binding term explicitly, and the clutch unbinding and clutch extension terms implicitly.

**Clutches with catch bond properties enhance traction forces**

In this section, we proceed to explore the effect of catch bond adhesion properties on force transmission. We assume that both catch and slip bond dissociation kinetics depends exponentially on clutch force. Therefore, the conservation equation for the probability density now reads:

$$\frac{\partial P_b}{\partial t} = k_{on}(n_c - n_b)\delta(x_c) - \left(k_{off}^{slip} e^{\frac{\kappa_c |x_c|}{F_b^{slip}}} + k_{off}^{catch} e^{-\frac{\kappa_c |x_c|}{F_b^{catch}}}\right) P_b - v_e \frac{\partial P_b}{\partial x_c}, \tag{S86}$$

where $k_{off}^{slip}$ and $k_{off}^{catch}$ are the unloaded clutch slip and catch bond dissociation rates, respectively, and $F_b^{slip}$ and $F_b^{catch}$ are the characteristic forces of the slip and catch bonds, respectively. In dimensionless form, Eq. (S18) reads

$$\frac{\partial P_b^*}{\partial t^*} = \tau(1 + \theta n_b^*)(1 - n_b^*)\delta^*(x_c^*) - \left(e^{F|x_c^*|} + K_{off}\, e^{-\frac{F|x_c^*|}{\chi_b}}\right) P_b^* - v_e^* \frac{\partial P_b^*}{\partial x_c^*}, \tag{S87}$$

where we have introduced two additional dimensionless numbers:

$$K_{off} = \frac{k_{off}^{catch}}{k_{off}^{slip}}, \qquad \chi_b = \frac{F_b^{catch}}{F_b^{slip}} \tag{S88}$$

The dimensionless clutch strain rate $v_e^*$ and the dimensionless first moment of the probability density over clutch extensions $\ell_b^*$ appearing in Eq. (S19) have been defined in Eq. (15) and (16), respectively. The numerical solution of Eq. (S19) is shown in Fig. (S6). For low values of the parameter $\omega$, force transmission depends inversely on the parameter $\chi_b$ for the whole range of substrate stiffnesses, since the effective clutch dissociation rate at low clutch extensions scales with $\chi_b$ due to catch bond behavior. For high values of the parameter $\omega$, force transmission is only sensitive to changes in the parameter $\chi_b$ for protrusions on soft enough substrate stiffnesses. On sufficiently rigid substrates, clutch loading is fast, slip bonds break by force, and the protrusion undergoes frictional slippage. Interestingly, the optimum substrate stiffness for maximal force transmission is not sensitive to the changes in the catch bond force parameter $\chi_b$, as shown in Fig. (S6).
